# Unbiased metastatic niche-labeling identifies estrogen receptor-positive macrophages as a barrier of T cell infiltration during bone colonization

**DOI:** 10.1101/2024.05.07.593016

**Authors:** Zhan Xu, Fengshuo Liu, Yunfeng Ding, Tianhong Pan, Yi-Hsuan Wu, Jun Liu, Igor L. Bado, Weijie Zhang, Ling Wu, Yang Gao, Xiaoxin Hao, Liqun Yu, David G. Edwards, Hilda L. Chan, Sergio Aguirre, Michael Warren Dieffenbach, Elina Chen, Yichao Shen, Dane Hoffman, Luis Becerra Dominguez, Charlotte Helena Rivas, Xiang Chen, Hai Wang, Zbigniew Gugala, Robert L. Satcher, Xiang H.-F. Zhang

**Affiliations:** Lester and Sue Smith Breast Center, Baylor College of Medicine, One Baylor Plaza, Houston, TX 77030, USA; Dan L. Duncan Cancer Center, Baylor College of Medicine, One Baylor Plaza, Houston, TX 77030, USA; Department of Molecular and Cellular Biology, Baylor College of Medicine, One Baylor Plaza, Houston, TX 77030, USA; Medical Scientist Training Program, Baylor College of Medicine, Houston, TX 77030, USA; Graduate Program in Cancer and Cell Biology, Baylor College of Medicine, One Baylor Plaza, Houston, TX 77030, USA; Graduate Program in Integrative Molecular and Biomedical Sciences, Baylor College of Medicine, One Baylor Plaza, Houston, TX 77030, USA; Graduate Program in Immunology and Microbiology, Baylor College of Medicine, One Baylor Plaza, Houston, TX 77030, USA; Graduate Program in Development, Disease Models, and Therapeutics, Baylor College of Medicine, One Baylor Plaza, Houston TX 77030, USA; Department of Orthopedic Oncology, University of Texas, MD Anderson Cancer Center, Houston, TX, United States of America; Department of Orthopedic Surgery and Rehabilitation, University of Texas Medical Branch, Galveston, TX, USA; McNair Medical Institute, Baylor College of Medicine, One Baylor Plaza, Houston, TX 77030, USA; College of Natural Sciences, University of Texas at Austin, 110 Inner Campus Drive, Austin, TX 78706, USA; Zhejiang Provincial Key Laboratory of Cancer Molecular Cell Biology, Life Sciences Institute, Zhejiang University, Hangzhou, Zhejiang 310058, China; Department of Orthopaedic Surgery, Second Affiliated Hospital, School of Medicine, Zhejiang University, Hangzhou, Zhejiang 310058, China

## Abstract

Microenvironment niches determine cellular fates of metastatic cancer cells. However, robust and unbiased approaches to identify niche components and their molecular profiles are lacking. We established Sortase A-Based Microenvironment Niche Tagging (SAMENT), which selectively labels cells encountered by cancer cells during metastatic colonization. SAMENT was applied to multiple cancer models colonizing the same organ and the same cancer to different organs. Common metastatic niche features include macrophage enrichment and T cell depletion. Macrophage niches are phenotypically diverse between different organs. In bone, macrophages express the estrogen receptor alpha (ERα) and exhibit active ERα signaling in male and female hosts. Conditional knockout of *Esr1* in macrophages significantly retarded bone colonization by allowing T cell infiltration. ERα expression was also discovered in human bone metastases of both genders. Collectively, we identified a unique population of ERα+ macrophages in the metastatic niche and functionally tied ERα signaling in macrophages to T cell exclusion during metastatic colonization.

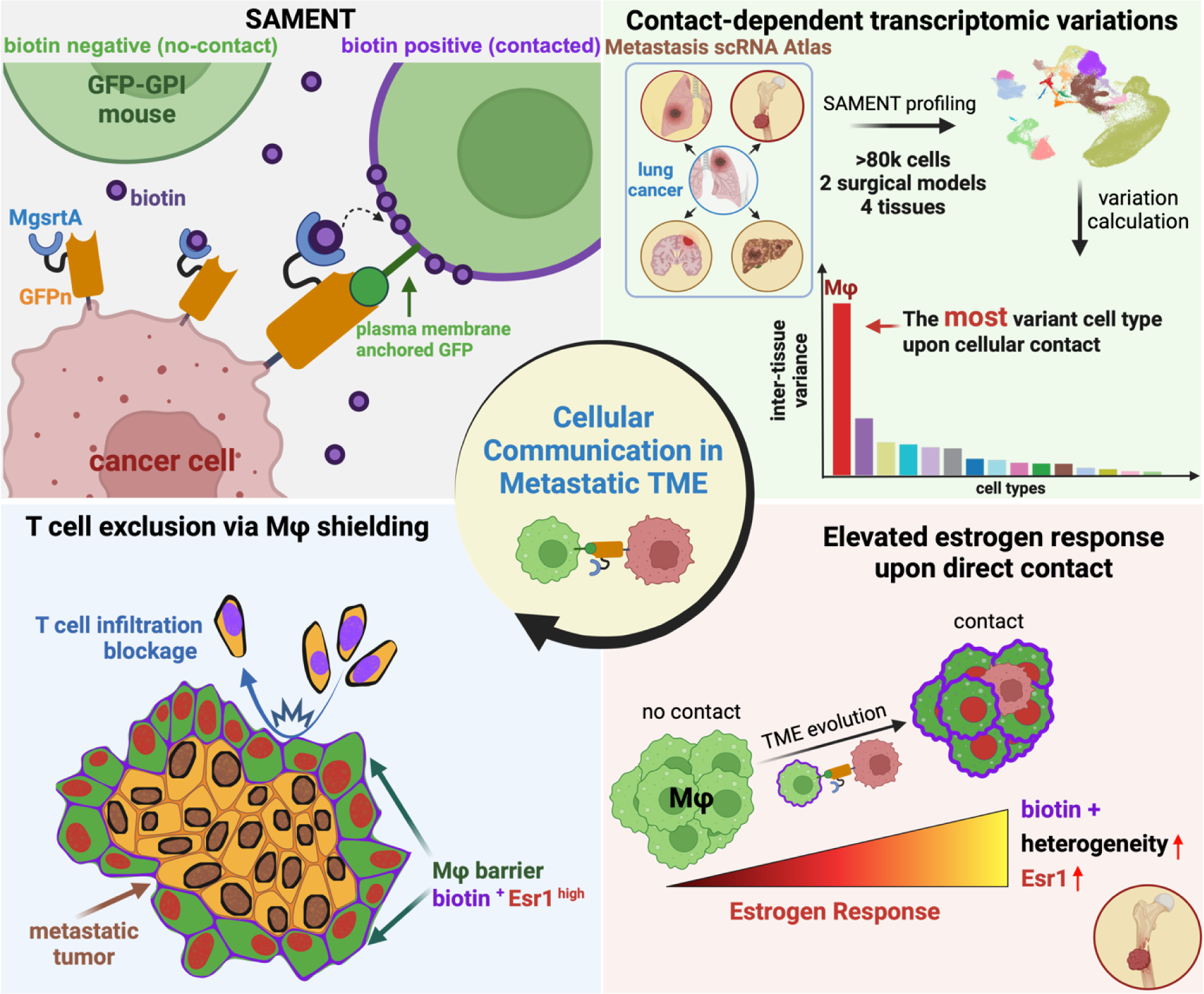

**HIGHLIGHTS:**

- SAMENT is a robust metastatic niche-labeling approach amenable to single-cell omics.
- Metastatic niches are typically enriched with macrophages and depleted of T cells.
- Direct interaction with cancer cells induces ERα expression in niche macrophages.
- Knockout of *Esr1* in macrophages allows T cell infiltration and retards bone colonization.

## INTRODUCTION

Throughout tumor progression and evolution, cancer cells are constantly interacting with the host, and this interaction profoundly impacts their cellular fates, metastatic behaviors and therapeutic responses. Numerous previous studies elucidated the roles of specific microenvironment niches (i.e., cells that are immediately adjacent to cancer cells) to the progression of metastasis. For instance, dormant disseminated tumor cells (DTCs) in the bone marrow tend to remain adjacent to the perivascular niche^1,2^. Endothelial cells are shown to induce mesenchymal-to-epithelial transition of DTCs^3^. Bone remodeling may activate DTCs and attract them to the osteogenic niche comprised of osteoblasts and their precursors^4^. Osteogenic cells and cancer cells can form heterotypic adherens junctions (hAJs) and gap junctions (GJs), which in turn activate the mTOR and calcium signaling, respectively, to promote progression of micrometastases^5–7^. Interaction with the osteogenic cells transiently reduces estrogen receptor alpha (ERα) expression and increases stem cell properties in the cancer cells^8^, which may in turn promote bone colonization, therapeutic resistance, and even further dissemination to other organs^9^. Taken together, the crosstalk between cancer cells and different BME niches appear to be of pivotal importance. However, this crosstalk is usually difficult to investigate because of its dynamic nature^10^ – many interactions may be transient and occur simultaneously between cancer cells and multiple other cell types. Thus, an unbiased approach is highly critical to capture all niche cells at specific stages of tumor progression.

Sortase A (SrtA) is a bacterial enzyme linking proteins to cell walls. It catalyzes cleavage of the threonine of the LPXTG motif, followed by fusion of this cleaved peptide to another protein with (Glycine)_n_ at the opposite end^11^. Recently, SrtA was adopted to study cell-cell interactions mediated by tight receptor-ligand interactions^12^. In these studies, one cell type was engineered to express SrtA-conjugated ligands. Upon provision of LPXTG-tag substrates, SrtA could transfer tag-LPXTG to the cognate receptors expressed on the other cell type. This strategy is powerful but is limited to only studying one ligand-receptor pair at a time. In this study we reported a novel system named SrtA-based MicroEnvironment Niche Tagging (SAMENT). By combining SrtA and synthetic ligand-receptor binding, we aim to label any cells that are physically encountered by cancer cells. A previous method was based on expression of lipid permeable mCherry (sLP-mCherry)^13^. Compared to sLP-mCherry, labeling by SAMENT is dependent on simultaneous induction of sortase and provision of substrates, and is amenable to downstream single-cell omics and spatial transcriptomic analyses. The robustness and specificity of SAMENT enabled us to conduct cellular and molecular profiling of the tumor immune microenvironment (TIME) landscape of different cancer models, in different organs and temporal stages, and under different treatments – thereby establishing a comprehensive metastatic niche atlas.

ER signaling is a major therapeutic target and biomarker of ER+ breast cancers^14^. Anti-ER agents, including selective estrogen receptor modulators (SERMs), aromatase inhibitors (Ais), and selective estrogen receptor degraders (SERDs), have been used in treating ER+ tumors in preventive, neoadjuvant, adjuvant and metastatic settings^15,16^. However, the roles of ER in non-cancer cells have just begun to be elucidated^17^. Through SAMENT, we observed enrichment of ERα+ macrophages in multiple cancer models in an unbiased fashion, and then performed conditional genetic knockout of *Esr1* in macrophages to investigate its functional contribution to bone colonization. Interestingly, ERα expression and signaling were also observed in human bone metastases of multiple cancer types. This discovery exemplified the utility of SAMENT and also raised an interesting possibility to target macrophage-derived ERα signaling in treating bone metastasis of multiple cancer types.

## RESULTS

### Design and validation of SAMENT

In cancer cells, a transgene consisting of fused the N-terminal mono-glycine recognition SrtA variant (mgSrtA)^18^ and a GFP nanobody (GFPn)^19^ are expressed under the control of a doxycycline-inducible promoter. MgSrtA is a variant of SrtA that recognizes one single glycine at the end of proteins such that a substantial proportion of cell surface proteins can serve as targets for unbiased labeling. GPI-GFP mice are used as hosts. These mice constitutively express outward-facing GFP on the surface of all cell types^20^ and bind GFPn, thereby allowing an unbiased tagging of any cells that come to the vicinity of cancer cells. When the cancer cells are provided with doxycycline (to induce mgSrtA-GFPn) and with biotin-labeled LPETGS (substrates of sortase), the GPI-GFP – GFPn binging stabilizes cell-cell interactions and allows mgSrtA to robustly label these adjacent cells for further analysis (**Figure 1A**).

**Figure 1.**
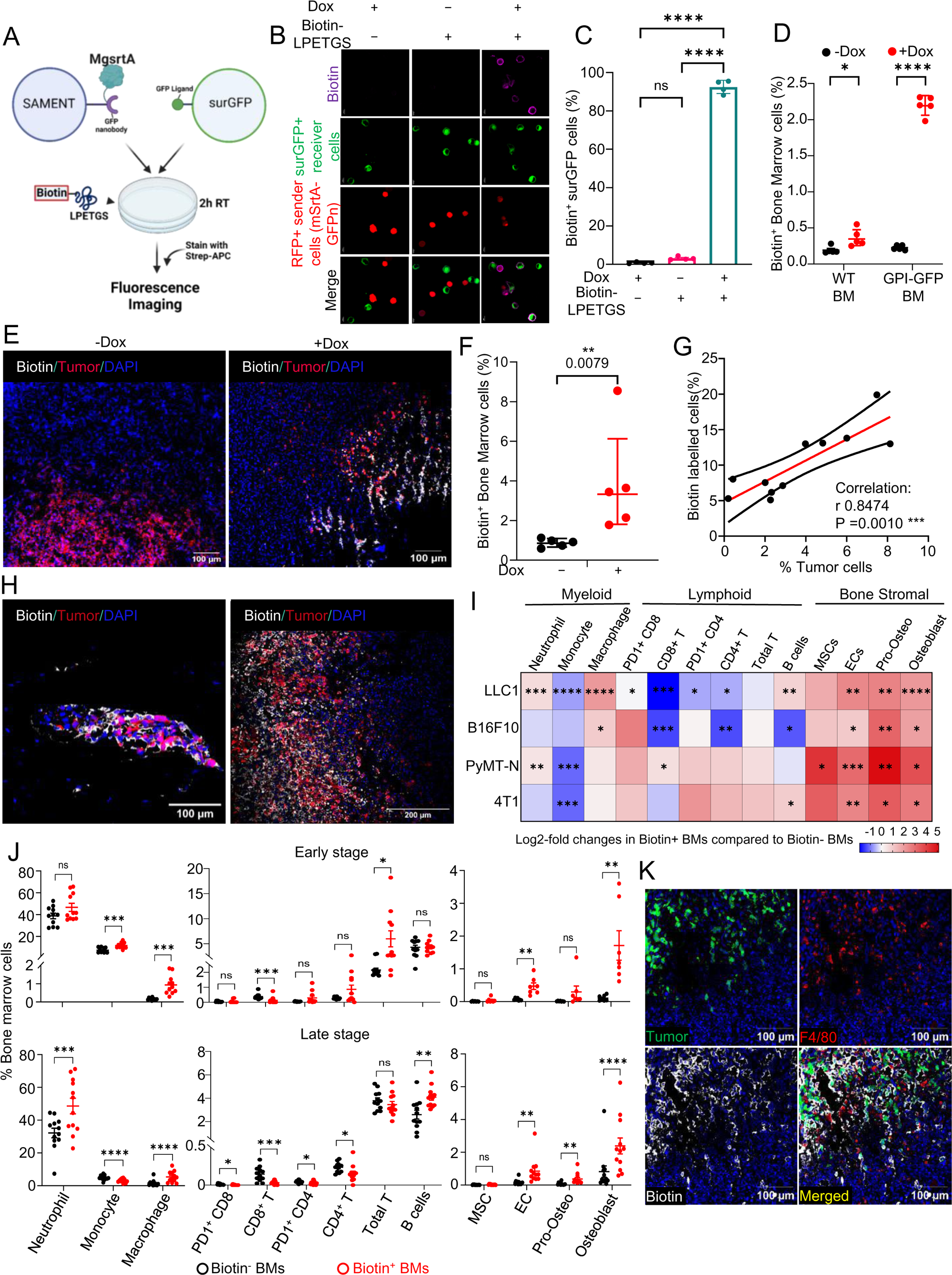
SAMENT approach enables specific labeling of metastatic niches. **(A)** Schematic representation of SAMENT cell labeling for surface-presented GFP (surGFP) cells *in vitro*. SAMENT+ sender cells express MgsrtA-GFPnanobody fusion protein on their surface, while surGFP+ receiver cells also present cell surface-bound GFP. Following a 2-hour co-culture at a 1:1 ratio and the addition of 500 uM biotin-LPETGS (SAMENT substrates), biotin-LPETGS transfers from SAMENT+ sender cells to surGFP+ receiver cells. **(B)** Fluorescence microscopy images depicting interaction-dependent Biotin labeling (purple) on the membrane of surGFP receiver cells (green), facilitated by SAMENT-functionalized LLC1 cells (red), exclusively in the presence of Doxycycline (inducing SAMENT expression) and biotin-LPETGS(SAMENT substrates). Data representative of at least two independent experiments. **(C)** Representative flow cytometric analysis of biotinylation of surGFP sender cells as shown in (B). Four biological replicates per group were tested. **(D)** Flow cytometric analysis of biotin labeling on GPI-GFP+ bone marrow cells isolated from GPI-GFP mice or WT bone marrow cells from WT mice, co-cultured *ex vivo* with SAMENT+ LLC1 cells. Five biological replicates per group were tested. **(E)** Representative fluorescence microscopy images and flow cytometric analysis reveal efficient biotinylation of GPI-GFP+ bone marrow cells exclusively when SAMENT is expressed by cancer cells (LLC1) during doxycycline treatment. **(F)** Quantitation of images represented by (E). Control group: n=5 mice; Doxycycline group: n=5 mice. **(G)** Examining the correlation between the percentage of biotin-labeled niche cells and the percentage of SAMENT+ cancer cells in metastatic bone analyzed by FACS. Analysis with different cancer cell frequencies (n=11 mice). **(H)** Representative fluorescence images of SAMENT labeling in micro-metastases and macro-metastases *in vivo*. **(I)** Log2-fold changes illustrate the relative frequency of each cell type among biotin+ bone marrow cells compared to that among biotin-bone marrow cells of GPI-GFP mice across different tumor models. LLC1, n=12 mice; B16F10, n=7 mice; Pymt-N, n=8 mice; 4T1, n=7 mice. **(J)** Quantification of pivotal immune and stromal cell populations in biotin+ bone marrow, contrasted with their biotin-counterparts, during early-stage (10 days) (n=11, For osteogenic population, n=7 mice) and late-stage (3 weeks) (n=12 mice) bone metastasis models induced by IIA injection with LLC1 cells. **(K)** Fluorescence imaging captures SAMENT-mediated biotinylation of macrophages (F4/80+) in the LLC1 bone metastasis model. Data representative of two independent experiments. Data are presented as geometric mean ± geometric SD in C, D, F; as mean ± SEM in J. P values were assessed by repeat measure one-way ANOVA followed by least significant difference (LSD) test in C; by unpaired t-test in D; by Mann–Whitney test in F; by Pearson correlation, two tailed in G; by paired t-test in I, J.

We first tested the SAMENT system *in vitro*. Lewis Lung Carcinoma cells (LLC1) were engineered to express RFP/mgSrtA-GFPn or GFP expressed on cell surface (surGFP) to generate red sender cells or green receiver cells, respectively. The sender and receiver cells were then co-cultured. We observed that cell labeling between senders and receivers occurred if and only if the biotin-labeled LPETGS peptides and doxycycline are simultaneously provided (**Figure 1B, 1C** and **S1A-S1C**). Intracellular expression of GFP in receiver cells did not result in labeling (**Figure S1D-S1F**). Within a wide range, the fraction of labeled receiver cells (over all receiver cells) is linearly proportional to incubation time (0-4 hours) (**Figure S1G-S1H**), concentration of LPETGS peptides (0.01-0.5 mM) (**Figure S1I-S1J**), and doxycycline (0-1 µg/ml) (**Figure S1K**). When the substrate was washed off, the biotin signals remained unchanged for about four hours, and then decayed to the background level in 24 hours with a half-life of about 16 hours (**Figure S1L**).

We then tested bone marrow (BM) derived cells as receiver cells. These cells were extracted from GPI-GFP or control mice and co-cultured with sender cancer cells. As expected, labeling of host BM cells only occurred when 1) GPI-GFP is on receiver cells, 2) mgSrtA is induced by doxycycline, and 3) LPETGS substrate is provided (**Figure1D, S1M and S1N**). Taken together, these results support that SAMENT works according to our design *in vitro* and exhibits a strong regulatability and specificity.

We then performed *in vivo* validation of SAMENT. Given our previous knowledge of bone metastatic niches, we decided to use bone as a major model. Cancer cells expressing inducible mgSrtA-GFPn were delivered to bone through intra-iliac artery (IIA) injection^21^. Upon establishment of bone lesions, we examined biotin-tagging in animals with or without provision of doxycycline to induce mgSrtA-GFPn. It was evident that the labeling only occurred when the SAMENT system is induced by doxycycline (**Figure 1E**). Flow cytometry analyses showed that induction of SAMENT resulted in a 6-8 fold more biotin labeling compared to background level (**Figure 1F**), suggesting a ∼70% specificity. Moreover, the frequencies of biotin+ cells are proportional to tumor burdens (**Figure 1G**). Importantly, this labeling is effective in both microscopic and macroscopic lesions (**Figure 1H**). This data supported the effectiveness of SAMENT *in vivo*.

We then extended SAMENT to multiple models and determined if we could replicate previous findings. We sorted biotin+ vs. biotin-cells based on expression of surface markers using flow cytometry. The frequency of each cell type among biotin+ cells was normalized to that among biotin-cells to deduce enrichment or depletion in the metastatic niche. In all four models, we observed a consistent enrichment of endothelial cells and osteoblasts/osteoprogenitor cells (**Figure 1I**), which is consistent with previous discoveries of roles of perivascular niches^1–3^, osteogenic niches^5–7,22^, and skeletal stem cells^23^. In addition, we also noticed unbalanced distribution of monocytes, macrophages, and T cells between biotin+ vs. biotin-populations. In particular, metastatic niches appear to be devoid of T cells, which is expected according to the paradigm of immune editing. The simultaneous depletion of monocytes and enrichment of macrophages suggest accelerated monocyte-to-macrophage differentiation upon encountering cancer cells (**Figure 1I**).

The inducibility of SAMENT allowed us to examine different temporal stages of bone colonization. Comparison between an early time point (10 days after IIA injection) and a relatively late time point (21 days after IIA injection) revealed evolution of metastatic niche. The noticeable changes with time include recruitment of neutrophils, reduction of monocytes, further increase of macrophages, and exacerbated depletion of T cells (**Figure 1J**). The prominent changes in myeloid cells prompted us to validate these findings by multiplexed immunofluorescence staining. Indeed, both macrophages (F4/80+) and neutrophils (Ly6G) are enriched in the metastatic niche (**Figure 1K** and **S1O**).

### SAMENT reveals a molecular and cellular atlas of metastatic niches

The “seed-and-soil” hypothesis has been used to understand preferential metastatic colonization in different organs^24,25^. The crosstalk between disseminated cancer cells and the immediate neighboring cells constitute a significant part of “seed and soil” interactions. Having established SAMENT and validated it in bone metastasis, we went on to determine how the same cancer cells may encounter and adapt to various milieus when arriving at different organs. To this end, we ignored organ-specific cells (e.g., osteoblasts in bone and hepatocytes in liver) and only focused on immune cells to allow meaningful inter-organ comparisons.

First, we validated that SAMENT worked as expected in lung and liver (**Figure 2A**). We then performed intracardiac injection of LLC1 cells to obtain experimental metastases in multiple different organs. FACS analyses on biotin+ niche cells from bone, brain, lung and liver metastases are compared. Similar to bone, metastases from other organs are also devoid of T cells but relatively enriched with macrophages (**Figure 2B**). However, there are also interesting differences noticed among organs. For instance, liver metastases exhibited less depletion of T cells and there appears to be a decrease as opposed to increase of neutrophils that are directly contacting cancer cells in lung metastases. These differences warrant further examination using additional models and lay the foundation for future research.

**Figure 2.**
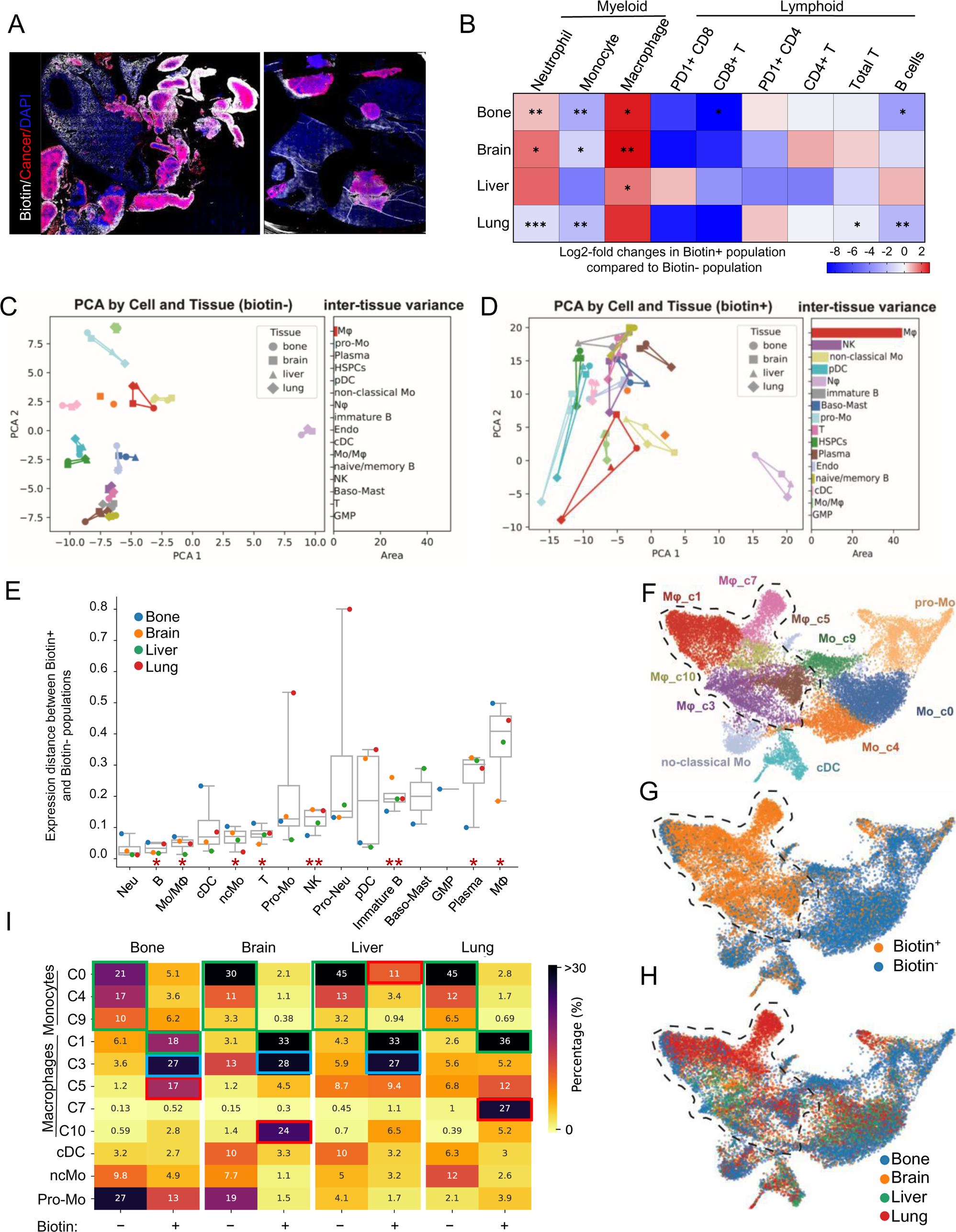
Cellular and molecular profiling of metastatic niches identify macrophages as a major component exhibiting phenotypic diversity and plasticity in different organs. **(A)** Fluorescence images depict SAMENT-mediated biotinylation of cells surrounding tumors in diverse tissues, including the lung (left) and liver (right). **(B)** Log2-fold changes depict alterations in the percentage of specific cell populations within biotin+ tissue population relative to corresponding biotin-tissue population across different organs with LLC1 tumor in GPI-GFP mice (n=5 mice). P values were assessed by paired t-test. **(C)** Principal components analysis (PCA) of biotin-niche cells across diverse tissues with LLC1 tumor. Lines connects tissue points for each cell type were drawn on the PCA plot, and the enclosed areas, reflecting inter-tissue transcriptomic variance for each cell type. **(D)** Same as (C) except for biotin+ niche cells across diverse tissues with LLC1 tumor. **(E)** Expression distance between the biotin labeling groups. Distance is based on the transforming the correlation between the biotin positive and negative cells, per cell type, per tissue. Larger distance indicates more average transcriptional differences between biotin positive and negative cell populations. Significance was calculated based on the mean differences include all four tissues, t-test applied. **(F)** UMAP of unbiased sub-clustering of myeloid populations. Numerically denoted subclusters of monocytes and macrophages, with macrophage populations highlighted by enclosed dotted lines. **(G)** The same UMAP as (F) except that distinct biotin statuses of macrophages and monocytes are projected on UMAP, indicating their direct interaction with cancer cells. **(H)** The same UMAP as (F) denoting tissue origin of myeloid cell subpopulations. **(I)** Myeloid sub-population frequencies by tissues of origin and biotin statuses. Sub-population frequencies were calculated per tissue per biotin group. Sub-population shared by all four tissues, three tissues, or one tissue are highlighted by green, blue, or red, respectively.

We next investigated transcriptomic variations of the same cell types at different metastatic sites. Single-cell RNA-seq data of major cell types (**Figure S2A**) were first divided into biotin-and biotin+ (non-niche vs. niche) and then subjected to principal component analysis (PCA). In this analysis, the distance between any two points or the area of a polygon connecting multiple points indicate the degree of dissimilarity among the underlying data. Among the biotin-cells, the points representing a same cell type in different organs clustered tightly together, indicating close similarity among the same cell type from different organs (**Figure 2C**). However, large inter-organ variations were observed among biotin+ cells for many cell types especially for macrophages (**Figure 2D**). The phenotypic diversity of macrophages has been intensively studied before^26,27^. Here, our data indicate that direct contact with cancer cells induce phenotypic heterogeneity of macrophages in different organs to the extent exceeding their normal diversity across tumor-free tissues. NK cells were ranked 2^nd^ in terms of their variations across different metastatic sites. An unbiased analysis of niche NK cells revealed multiple macrophage regulation pathways (**Figure S2B**), suggesting macrophages as a downstream effector of NK cells in metastatic niches.

We then asked how biotin+ and biotin-cells vary from one another in different organs. To this end, we calculated the “distance” between biotin+ and biotin-cells and compared the average distance of each cell type in four different organs (**Figure 2E**). Macrophages were again the top cell type in this comparison, indicating that their interaction with tumor cells causes the largest phenotypic alteration. In fact, among the four organs, macrophages in bone exhibit the largest difference between biotin+ and biotin-cells, prompting further investigation elaborated in later sections. Other noteworthy tumor-induced changes include immature B cells (**Figure S2C**) and neutrophil and monocyte progenitors in lung metastasis (**Figure S2D**). Further analyses revealed interesting cancer-induced transcriptomic shifts in these cell types in an organ-specific fashion. Among all cell types, granulocyte-monocyte precursors (GMPs) were included because of our special interest stemming from our previous research^28^. Since GMPs are confined to the bone marrow, we could not analyze them in other organs. Nevertheless, biotin+ and biotin-GMPs displayed a substantial transcriptomic difference (**Figure S2E**) that is consistent with our previous finding of tumor-entrained systemic reprogramming of hematopoietic stem and precursor cells including GMP ^28^.

Having observed the remarkable macrophage heterogeneity and plasticity upon direct contact with tumor cells, we performed detailed analyses on isolated scRNA-seq data of the monocyte lineage (**Figure 2F-2H**) and identified several clusters. Tabulation of frequencies of these clusters across different metastatic sites and biotin statuses led to interesting findings (**Figure 2I**). Compared to biotin+ cells, biotin-cells enrich monocytes (Clusters C0, C4 and C9) similarly across all metastatic sites. On the other hand, the macrophage clusters (C1, C3, C5, C7 and C10) are the dominant populations in biotin+ cells but exhibited largely diverse distributions across different metastatic sites. Specifically, C1 and C3 abound in multiple sites, whereas C5, C7 and C10 are predominantly enriched in one specific organ. Taken together, the data suggests that C1 and C3 are monocyte-derived macrophages whereas C5, C7 and C10 are tissue-specific macrophages recruited to metastases. Indeed, detailed examination of known markers of macrophages of different origins support this hypothesis (**Figure S2F**).

Taken together, we have established an immune cell atlas of metastatic niches using SAMENT. We identified cell types that are preferentially contacted by cancer cells, and that are commonly or differentially enriched in metastases at different sites. Among all cell types, macrophages occur most frequently surrounding disseminated cancer cells and appear to be phenotypically reprogrammed upon interaction with metastases. Therefore, we will focus on the roles of macrophages in metastatic niches for the remainder of this study.

### Macrophages in the metastatic niche express estrogen receptor alpha (ERα)

Differentially expressed genes and pathways were identified by unbiased comparison between biotin+ and biotin-macrophages across all tissues (**Figure 3A and 3B**). We noticed a number of up-regulated pathways are related to ERα signaling (**Figure 3B**). ER signaling is a major driver of ER+ breast cancer. It also plays an important role in many other cell types including macrophages, T cells, osteoblasts and osteoclasts. The immunoregulatory functions of ER signaling may result from evolutionary needs for tolerating fetus during pregnancy. The immunosuppressive impact of ERα in macrophages was recently reported in melanoma ^29–31^. The pleiotropy of ERα makes it an interesting target both for validation of SAMENT methodology and for discovery of novel biology in metastasis. We verified the expression of ERα in macrophages adjacent to metastases by immunofluorescence staining (**Figure 3C**). In addition, among other immune cell types macrophages exhibit relatively high expression of ERα (**Figure S3A**).

**Figure 3.**
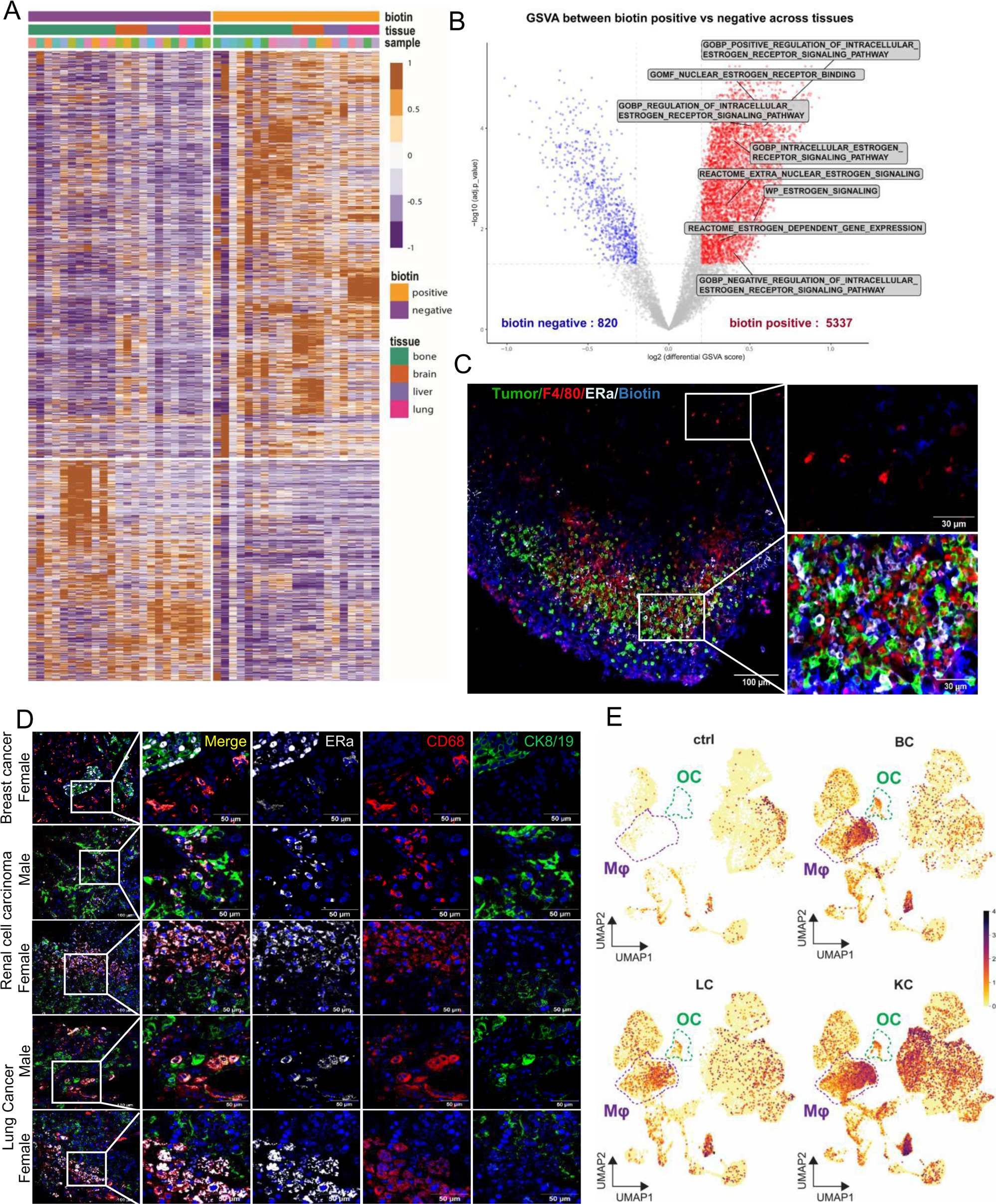
Macrophages within the bone metastasis niche exhibit heightened ERα signaling. **(A)** Heatmap displays the significant differentially expressed genes (DEGs) between biotin+ and biotin-macrophages in metastasitic LLC1 tumor tissues. With rows representing genes and columns are samples. Both rows and columns are hierarchical clustered. **(B)** Volcano plot shows the differentially enriched pathways between biotin+ and biotin-niche macrophages, include all organs with LLC1 tumor. Estrogen-related pathways across the entire MsigDB dataset are highlighted, exclusively enriched in biotin+ populations. **(C)** Fluorescence imaging reveals that within the bone tumor microenvironment, F4/80^+^ murine macrophages labeled by SAMENT^+^ tumor cells with surface biotinylation exhibit higher ERα expression compared to those located distantly from the tumor area. Data representative of two independent experiments. **(D)** Representative immunofluorescence staining reveals Cytokeratin 8/19-positive cancer cells in bone metastases from multiple cancer types exhibit co-localization of ERα and CD68 (human macrophage marker) in both male and female patients. Data representative of at least three independent patient samples. **(E)** Normalized ESR1 Expression Levels in Human Bone Metastasis of Breast Cancer (BC), Lung Cancer (LC) and Kidney Cancer (KC) as visualized by Feature Plot on UMAP Coordinates. The plot highlights the macrophage and osteoclast cell populations where ESR1 expression is evident.

We next asked if ERα expression in macrophages also occurs in human bone metastases. In a companion study, we collected and analyzed clinical 34 samples of human bone metastases stem from a variety of primary cancer sites. By immunofluorescence staining, we observed robust macrophage ERα expression in 10 metastases including male patients (**Figure 3D and S3B**). For comparison, we also examined a primary colorectal cancer and did not observe ERα expression in associate macrophages (last column of Figure S3B). Enhanced expression of ESR1 in macrophages was further validated by single-cell RNA sequencing in human bone metastases from breast cancer (BC), lung cancer (LC), and kidney cancer (KC), compared to healthy bone marrow (**Figure 3E and S3C**). Additionally, osteoclasts, which are specialized tissue-resident macrophages in bone^32^, were also found to exhibit increased ESR1 expression. Detailed analyses of these bone metastasis samples were conducted in our companion study.

### ERα expression is caused by direct contact with cancer cells

Two potential mechanisms may drive the overexpression of ERα in biotin+ macrophages: 1) recruitment and enrichment of pre-existing ERα+ macrophages and 2) upregulation of ERα expression when macrophages encounter cancer cells. To distinguish these two possibilities, we performed *in vitro* co-culture experiments using cancer cells and bone marrow-differentiated macrophages expressing luciferase/GFP reporters driven by estrogen responsive elements (EREs). Compared to the monoculture control, both luciferase activities and GFP expression were significantly elevated in co-cultures (**Figure 4A**), suggesting up-regulation of ERα activities upon interaction with cancer cells. This experiment was repeated in multiple cancer cell lines and the same observation was made in five out of seven lines (**Figure 4B**). We further verified that the upregulation of ERE activities is dependent on direct cell-cell contact (**Figure 4C**) and occurred through transcriptional increase of *Esr1* gene encoding ERα (**Figure 4D**). Finally, immunofluorescence staining of bone metastasis *in vivo* revealed selectively enhanced ERE activities in macrophages adjacent to tumor cells (**Figure S4A**).

**Figure 4.**
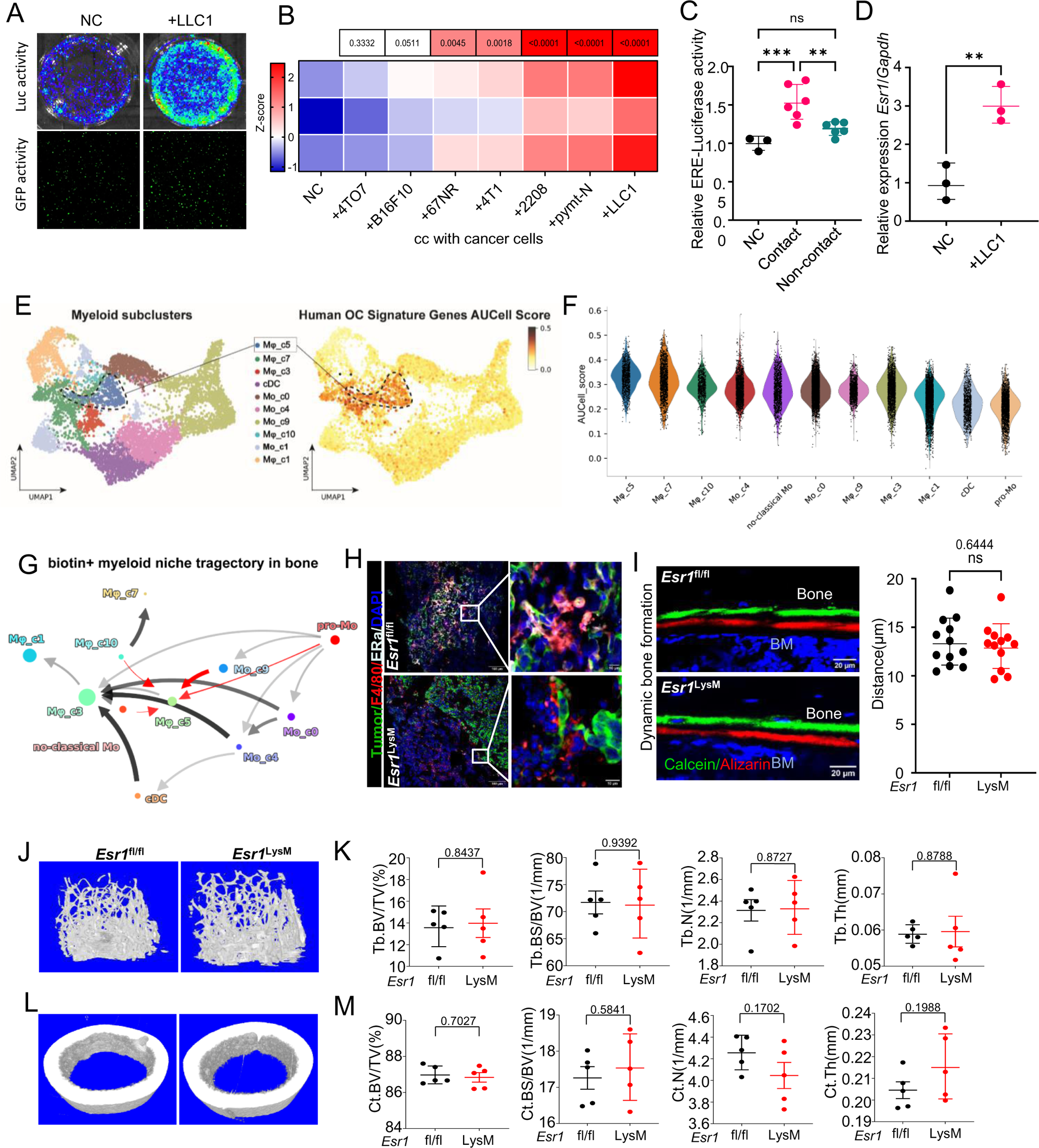
ERα expression in macrophages results from direct interaction with tumor cells and does not regulate normal bone homeostasis. **(A)** The representative bioluminescence image and fluorescence images reveal heightened luciferase and GFP expression in bone marrow-differentiated macrophages transfected with the ERE(Estrogen Response Elements)-Luc-T2A-GFP reporter following co-culture with LLC1 cells. Data representative of two independent experiments. **(B)** Heatmap shows that estrogen receptor transactivation was assessed through luciferase activity in bone marrow-differentiated macrophages transfected with the ERE-luciferase reporter and co-cultured with various types of cancer cells for 2 days. The luciferase data of the co-culture group were normalized to the mean value of the macrophage monoculture group (NC), and then transformed to Z-score. Three biological replicates per group were shown. **(C)** The relative ERE-luciferase activity of bone marrow-differentiated macrophages after a 2-day direct or indirect co-culture with LLC1 cells, normalized to macrophage culture (NC) only. **(D)** qPCR analysis of the expression of *Esr1* mRNA in bone marrow-differentiated macrophages only or co-culture with LLC1 cells. The expression was normalized to the mean value of the macrophage culture only (NC) group. Three biological replicates per group were shown. **(E)** Osteoclast identification in SAMENT single-cell dataset. UMAP shows unbiased subclustering of all myeloid cells (left). The expression of human OC signature markers were scored and projected on the UMAP (right). **(F)** Estrogen responsive pathway activity scored across the myeloid subclusters (E). Gene sets come from MsigDB database hallmark pathway terms “Hallmarks Estrogen Response Early” and “Hallmarks Estrogen Response Late”. **(G)** Trajectory inference for the myeloid subclusters in biotin positive populations. Osteoclasts (Mφ_c5) is one of the terminal states across the trajectory an it has three precursors: Mφ_c3, Mφ_c10 and Mo_c9. Red arrows indicate the estimated developmental trajectory. Thickness of arrows indicates frequencies of differentiation events. **(H)** A representative set of immunofluorescence images depicting ERα expression in tumor-adjacent murine macrophages(F4/80^+^) derived from *Esr1*^fl/fl^ and LysM-cre *Esr1*^fl/fl^ bone marrow. Data representative of two independent experiments. **(I)** Representative confocal images (left) and quantified rates (right) depicting new bone formation in *Esr1*^fl/fl^(n=3 mice) and LysM-cre *Esr1*^fl/fl^ mice (n=3 mice). Four independent images were measured for each mouse and included in Figure I. **(J)** Representative three-dimensional μ-CT images of trabecular bone of distal femurs isolated from 8-week-old female mice with *Esr1*^fl/fl^and LysM-cre *Esr1*^fl/fl^ genotypes. **(K)** Quantification of trabecular bones shown in (J) volume per tissue volume (Tb.BV/TV). Trabecular bone surface per bone volume (Tb.BS/BV). Trabecular number (Tb.N). Trabecular thickness (Tb.Th). (n=5 mice each group) **(L)** Representative three-dimensional μ-CT images of cortical bone of middle shaft of femurs isolated from 8-week-old female mice with *Esr1*^fl/fl^ and LysM-cre *Esr1*^fl/fl^ genotypes. **(M)** Quantification of cortical bones shown in (L) volume per tissue volume (Ct.BV/TV). Cortical bone surface per bone volume (Ct.BS/BV). Cortical number (Ct.N). Cortical thickness (Ct.Th). (n=5 mice each group) Data are presented as geometric mean ± geometric SD in C, D, I, K, M. P values were assessed by repeat measure one-way ANOVA followed by least significant difference (LSD) test in B, C; by unpaired t-test in D, I, K, M.

### ERα in monocytes and macrophages does not regulate bone formation or remodeling

Osteoclasts drive the osteolytic vicious cycle, which is a major mechanism underlying advanced bone metastases of multiple cancer types^33–36^. Monocytes and macrophages are precursors of osteoclasts. However, the osteoclast differentiation is determined by the microenvironment and intrinsic properties of precursors, and only occurs to a subset of monocytes and macrophages^30,31^. It is important to ask if ERα+ macrophages are indeed osteoclasts or their precursors. To address this question, we first identified osteoclasts among all myeloid cells by applying a set of osteoclast signatures^37^. Cluster C5 is uniquely enriched in bone metastasis (**Figure 2I**) and overexpresses the osteoclast signature (**Figure 4E)**, and was therefore identified as osteoclasts. ERα activity in C5 is heterogeneous, indicating that there are indeed ERα^high^ osteoclasts (**Figure 4F**). We then performed pseudotime analyses to deduce precursors of C5 and identified C10 (macrophages) and C9 (monocytes) (**Figure 4G**). Since C10 is predominantly found in brain metastases (**Figure 2I**), C9 is more likely the osteoclast precursors, and the ERα activity in this subset is low. Therefore, although some osteoclasts are ERα^high^, the major osteoclast precursors are ERα^low^.

Since a subset of osteoclasts exhibit ERα activity, we asked if blocking ERα signaling may affect bone formation or remodeling. To this end, we generated *Esr1*^LyzM^ mice with LyzM-cre driven *Esr1* knockout in monocytes, mature macrophages and granulocytes^38^, which include osteoclast precursors and differentiated osteoclasts. The knockout was validated by immunofluorescence staining (**Figure 4H**) and Western blots (**Figure S4B**). Loss of ERα in these cells did not generate discernible effects on bone formation (**Figure 4I**), nor did it affect trabecular or cortex bones (**Figure 4J-4M**). This is consistent with the fact that ER upregulation in macrophages is induced by direct contact with cancer cells but remains at a relatively low level in normal bones. Thus, perturbation of macrophage-derived ERα does not appear to alter bone homeostasis.

### Conditional knockout of *Esr1* in macrophages hinders metastatic colonization in bone

To examine if macrophage-derived ERα plays a role in bone metastasis, we performed IIA injection to introduce experimental bone metastases of LLC1, E0771, 4T1 and PyMT-M cells in syngeneic hosts with or without LysM-cre driven *Esr1* knockout. Since LLC1 is a lung cancer cell line and the known functions of ERα are gender-specific, we also investigated bone colonization of LLC1 in male vs. female hosts (**Figure 5A**). In all experiments, bone colonization in *Esr1*^LyzM^ animals was significantly retarded. Interestingly, for the LLC1 model, the impact of conditional *Esr1* knockout was even more striking in male animals (**Figure 5A**). This is consistent with patient data in **Figure 3D**, which clearly showed a similar level of ERα in bone metastases of male patients. Importantly, the decrease of metastatic burden was not only evident in the hind limbs subjected to IIA injection (**Figure 5B**), but in most models also noticeable in contralateral limbs (non-injected) (**Figure 5C**) and lungs (**Figure 5D**). These latter metastatic lesions resulted from further dissemination of the initial bone metastases as shown in our previous studies^9^.

**Figure 5.**
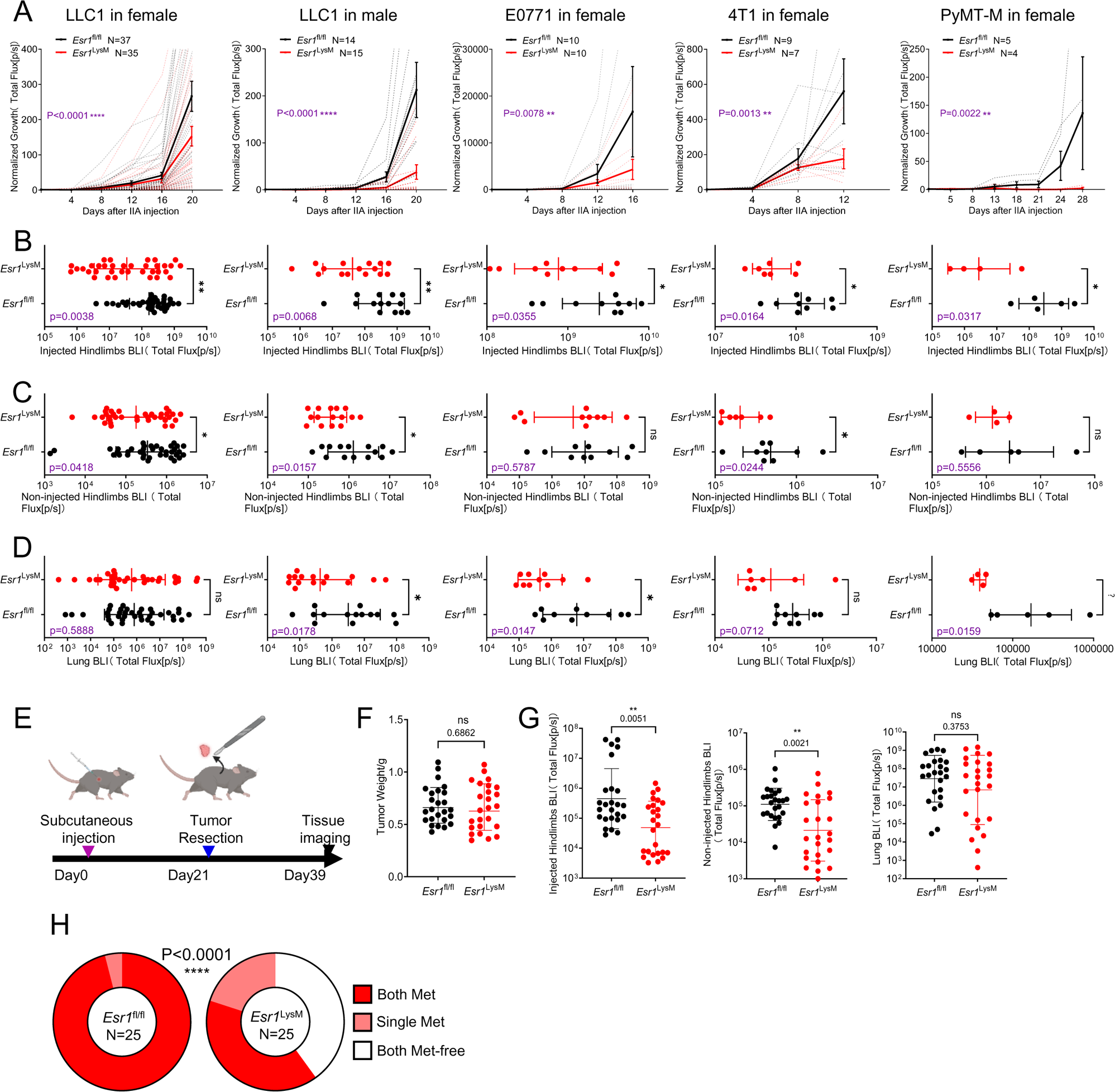
Conditional knockout of *Esr1* in macrophages significantly retarded tumor colonization in bone. **(A)** Normalized tumor growth curves of LysM-Cre *Esr1*^fl/fl^mice (n=35, 15, 10, 7, 4 mice) and their respective control mice (n=37, 14, 10, 9, 5 mice) following IIA injection with LLC1, E0771, 4T1, and PyMT-M breast cancer cells are depicted. The gender of the mice utilized in each experiment is indicated at the top of each respective figure. Each dotted curve represents an individual animal, whereas the highlighted curve shows the mean growth for each group in A. **(B)** Quantified *ex vivo* bioluminescence imaging intensities in hindlimb bones on the same side of IIA injection (Injected) of LysM-Cre *Esr1*^fl/fl^mice (n=35, 15, 10, 7, 4 mice) and their respective control mice (n=37, 14, 10, 9, 5 mice) are presented following IIA injection with LLC1, E0771, 4T1, and PyMT-M breast cancer cells. **(C)** Same as (B) except that the bioluminescence signals of hind limbs contralateral to IIA injection (non-injected) are quantitated. Models and sample sizes are indicated in (B). **(D)** Same as (B) except that the bioluminescence signals of lungs are quantitated. Models and sample sizes are indicated in (B). **(E)** Schematic of spontaneous bone metastasis assy. **(F)** Quantification of source subcutaneous tumor weight (n=25 mice in each group). **(G)** Quantification of spontaneous metastases (bioluminescence intensity [BLI]) to hind limb bones and lungs (n=25 mice in each group). **(H)** The ratio of detectable metastasis in both hindlimb bones in female LysM-Cre *Esr1*^fl/fl^ mice (n=25 mice) and their corresponding control mice (n=25 mice) injected with LLC1 lung carcinoma cells. Data are presented as mean ± SEM in A; geometric mean ± geometric SD in B, C, D, F, G. P values were assessed by Fisher least significant difference test post repeat measure two-way ANOVA test in A; by Mann–Whitney test in B, C, D, F, G and by Chi-square test in H.

LLC1 cells spontaneously metastasize to bone, which allows us to examine the role of macrophage ERα in a complete metastatic cascade (**Figure 5E**). Subcutaneous source tumors grew at a similar rate in *Esr1*^LyzM^ and control *Esr1*^fl/fl^ animals (**Figure 5F**), and were removed after reaching 1 cm^3^ (**Figure 5E**). 18 days after tumor resection, we harvested hind limb bones and lungs to assess metastases to these organs. A 5-to 10-fold difference was observed for both hind limbs (**Figure 5G-5H**). Lung metastases trended toward a reduction but did not reach statistical significance (**Figure 5G**).

To determine if the role of ERα is specific to macrophages, we also conditionally knocked out ERα by crossing *Esr1*^fl/fl^ with a number of other cre-lines that target major cell types in bone including Cx3cr1-cre, Osterix-cre, Col1a1-cre, S100A8-cre, and Itgax-cre in both experimental metastasis and spontaneous metastasis assays (**Figure S5**). Among these, *Esr1*^Cx3cr1^ animals exhibited significant decrease of bone colonization, albeit to a lesser degree compared to *Esr1*^LyzM^ (**Figure S5A-S5C**). CX3CR1+ marks a subset of macrophages^39^, and this result supports the role of ERα in macrophages. On the other hand, S100A8 is predominantly expressed in granulocytes ^40^, and Itgax (a.k.a. CD11c) is a dendritic cell marker. The lack of effect in these models distinguish macrophages from other major myeloid cell populations. Taken together, our results strongly support the hypothesis that ERα in macrophages plays an important role in bone colonization.

### Conditional knockout of *Esr1* in macrophages leads to increased T cell infiltration and activation in metastatic niches

We next set out to determine the cellular mechanism underlying the observed decrease of bone colonization upon *Esr1* knockout in macrophages. Major immune cell types in metastasis-associated bone microenvironment were quantitated using flow cytometry and compared between *Esr1*^LyzM^ and *Esr1*^fl/fl^ hosts but did not uncover significant differences (**Figure S6A and S6B**). Therefore, loss of ERα in macrophages does not seem to affect immune cell frequencies in bone metastases. This is contrasted by anti-PD1 treatment, which significantly increases CD8+ T cells (**Figure 6A**). However, multiplexed immunofluorescence staining revealed dramatic changes in the spatial distribution of T cells. In wild type hosts, T cells appear to be sequestered by ERα+ macrophages at the edge of metastatic lesions (**Figure 6B-6E**). This is consistent with the lack of T cells in biotin+ cells in SAMENT experiments (**Figure 1I**), which indicates exclusion of T cells and scarcity of direct cancer-T cell interactions. When ERα is depleted in macrophages, metastasis-infiltrating, Granzyme B^+^ T cells significantly increased (**Figure 6D-6E**). Thus, conditional knockout of *Esr1* in macrophages appears to impact T cells in a different fashion as compared to anti-PD1 treatment.

**Figure 6.**
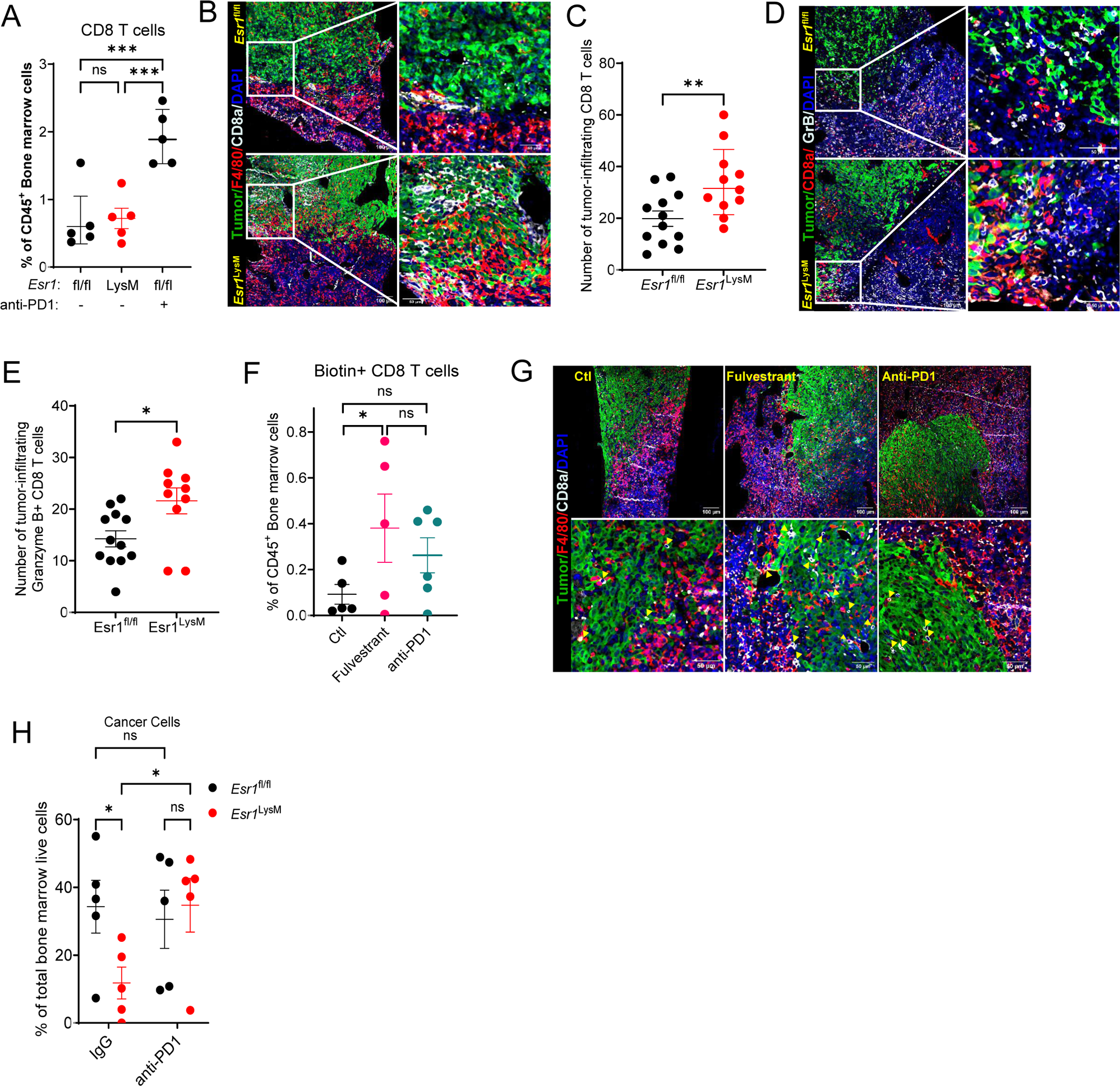
ERα signaling in macrophages excludes T cells from metastatic lesions. **(A)** CD8 T cell quantification was conducted in the hindlimb bones of mice injected with LLC1 cells. This included *Esr1*^fl/fl^ mice with and without anti-PD1 treatment (n=5 mice each), as well as LysM-Cre *Esr1*^fl/fl^ mice (n=5 mice). The frequency of CD8 T cells remained unaffected by *Esr1* knockout; however, it significantly increased following anti-PD1 treatment. **(B)** A representative series of immunofluorescence images illustrates the heightened infiltration of CD8 T cells into the tumor region in bone metastatic tumors of LysM-Cre *Esr1*^fl/fl^ mice compared to control mice. In the images, CD8 T cells are marked in white, F4/80+ macrophages in red, and LLC1 tumor cells in green. **(C)** Quantification of tumor-infiltrating CD8 T cells in LysM-Cre *Esr1*^fl/fl^ mice compared to their control *Esr1*^fl/fl^ counterparts. **(D)** A representative series of immunofluorescence images illustrates the enhanced infiltration of Granzyme B+ CD8 T cells into the tumor region in bone metastatic tumors of LysM-Cre *Esr1*^fl/fl^ mice compared to control mice. In the images, CD8 T cells are marked in red, Granzyme B in white, and LLC1 tumor cells in green. **(E)** Quantification of tumor-infiltrating Granzyme B+ CD8 T cells in LysM-Cre *Esr1*^fl/fl^ mice compared to their control *Esr1*^fl/fl^ counterparts. **(F)** Quantification of biotin+ CD8 T cells was carried out in the hindlimb bones of mice treated with either fulvestrant or anti-PD1 following IIA injection with SAMENT+ LLC1 cells. The findings suggest that compared to anti-PD1 treatment, fulvestrant enhances the infiltration of CD8 T cells into metastatic tumors. Control group: n=5 mice; Fulvestrant group: n=5 mice; anti-PD1 group: n=5 mice. **(G)** A series of representative immunofluorescence images vividly depict the heightened infiltration of CD8 T cells into the tumor region within bone metastatic tumors of mice subjected to fulvestrant treatment. In these images, CD8 T cells are highlighted in white, F4/80+ macrophages in red, and LLC1 tumor cells in green. The infiltrated CD8 T cells are indicated by yellow arrows. **(H)** Cancer cell quantification was performed in the hindlimb bones of mice injected with LLC1 cells, including LysM-Cre *Esr1*^fl/fl^ mice (n=5 mice) and their control *Esr1*^fl/fl^ counterparts (n=5 mice), with or without anti-PD1 treatment, following IIA injection. Data are presented as geometric mean ± geometric SD in A, C; mean ± SEM in E, F, H. P values were assessed by repeat measure one-way ANOVA followed by least significant difference (LSD) test in A; by unpaired t-test in C, E, F; by Fisher least significant difference test post repeat measure two-way ANOVA test in H.

We next asked if fulvestrant, a SERD clinically used to treat metastatic breast cancer, could achieve the same effect as *Esr1* knockdown. Indeed, both SAMENT (**Figure 6F**) and immunofluorescence staining (**Figure 6G**) confirmed that fulvestrant, but not anti-PD1, significantly increases T cell infiltration. Interestingly, we observed decreased tumor burden in *Esr1*^LyzM^ hosts but not in anti-PD1 treated animals (**Figure 6H**), suggesting that T cell infiltration is more important in controlling metastatic colonization. However, combining anti-PD1 and ER knockout did not achieve additive or synergistic effects (**Figure 6H**), suggesting more complicated interactions between ERα signaling in macrophages and the PD1 pathway in T cells. The underlying mechanism remains unclear and beyond the scope of this study, although it may be caused by increased monocytes and macrophages that selectively occur to animals subjected to the combined treatment (**Figure S6C**). Further studies are needed to identify other combination strategies to leverage the increased T cell infiltration when macrophage ERα is inhibited.

## DISCUSSION

We established SAMENT to unbiasedly profile microenvironment niche cells that are immediately adjacent to metastatic seeds. Approaches with similar goals were reported before, including expression of a secreted fluorescence mCherry containing a lipid-permeable transactivator of transcription (TATk) peptide (sLP-mCherry)^13^. Compared to these previous approaches, SAMENT is more tightly controlled as the labeling is simultaneously dependent on both substrate and the doxycycline inducer. Furthermore, the labeling only occurs to neighboring cells, and the spatial proximity is “locked” by GPI-GFP:GFPn interaction upon GFPn induction. These elements render SAMENT highly specific and efficient as validated by immunofluorescence staining. Further development of SAMENT may enable us to answer more specific questions. For instance, conditional expression of GPI-GFP in selective lineages of host cells will allow us to focus only on specific niche cells.

Macrophages play important roles in tumor progression including metastases at different sites and development of therapeutic resistance ^41–47^. Aside from being precursors of osteoclasts, the specific functions of macrophages in bone metastasis have only recently been studied in breast and prostate cancers^26,27^. Herein, our study elucidated a novel subset of macrophages expressing ERα. Interestingly, ERα expression is induced by direct cell-cell contact between cancer cells and macrophages, and conditional knockout of *Esr1* appears to alleviate the exclusion of T cells from infiltrating metastatic lesions. The immunosuppressive effects of the ERα pathway in macrophages have been investigated before^29^. However, the spatial relationships among ERα+ macrophages, cancer cells and T cells, as revealed by SAMENT and immunofluorescence staining, have never been discovered. This highlights the relevance of spatial information in our understanding of tumor-host interactions. Toward this end, SAMENT provides an objective measurement of geographic, immediate proximity.

Despite its prominent female-specific roles, ERα signaling is also involved in multiple physiological processes in males^48^. This has been largely neglected in cancer research. In this study, we were surprised to find that conditional knockout of *Esr1* generated an even stronger effect on bone metastases in male animals. Consistently, we also observed robust ERα expression in macrophages associated with bone metastases in male patients. These observations raised an interesting possibility of applying well established anti-ER endocrine therapies to male patients with bone metastases of multiple cancer types. The downstream effectors of ERα signaling in macrophages will need to be identified in future research.

The importance of ERα signaling in macrophages may be questioned in breast cancer, as endocrine therapies have been shown to have no benefit for ER-patients^49^. However, the previous studies have never focused exclusively on bone metastases, which are relatively infrequent and often co-occurring with metastases to other organs in ER-diseases. Furthermore, as shown in the final set of experiments, inhibition of ERα in macrophages may not be effective by itself but could synergize with immunotherapies because it facilitates T cell infiltration into metastatic lesions. Although we did not observe synergy between ERα knockout in macrophage and anti-PD1 treatment, it is still worth exploring the combinatory effects with other immunotherapies. Therefore, our findings may warrant future clinical trials on combined endocrine and immunotherapies on patients with bone metastases, and this combination may be extended to other cancer types and to patients of both genders.

## LIMITATIONS

SAMENT effectively labels niche cells because the tight interaction between GPI-GFP and GFPn helps stabilize cell-cell interactions. However, this also poses a limitation as the same interaction also hinders cancer cells from migrating. As such, we can only induce GFPn immediately before we terminate the experiment and examine a snapshot of metastatic niches. Complementary approaches are under development to track niche cells longitudinally even after they depart from metastatic cells.

## ACKNOWLEDGEMENTS

We are grateful to the discussion and suggestions from Zhang Laboratory members. We also thank Dr. Anna-Katerina Hadjantonakis (MSKCC) for providing the GPI-GFP mice, Dr. Michael P. Rout (Rockefeller University) for providing constructs of GFP nanobodies and Dr. Trey Westbrook (BCM) for providing constructs of pinducer20. X.H.-F.Z. is supported by US Department of Defense DAMD W81XWH-21-1-0790, HT9425-23-1-0493 NCI CA183878, NCI CA251950, NCI CA277838-01A1, NCI CA271498, NCI U01CA252553 and Breast Cancer Research Foundation. This project was supported by the Cytometry and Cell Sorting Core at BCM with funding from the CPRIT Core Facility Support Award (CPRIT-RP180672), the NIH (P30 CA125123, S10 RR024574, S10 OD025251); by the Pathology Core of Lester and Sue Smith Breast Center at BCM; by the Optical Imaging & Vital Microscopy (OiVM) core at BCM; by the Single Cell Genomics Core at BCM partially supported by NIH S10OD025240 and CPRIT RP200504; by the RNA In Situ Hybridization Core at BCM with funding from the PHS grant DK056338; by the Integrated Microscopy core at BCM with funding from NIH (DK56338, CA125123, ES030285), and CPRIT (RP150578, RP170719). We also acknowledge Joy Guo, Dr. Mira Jeong, Jason Kirk, Dr. Cecilia Ljungberg, Rena Mao and Joel M. Sederstrom at Baylor College of Medicine for their expert assistance.

## AUTHOR CONTRIBUTIONS

X.H.-F.Z. and Z.X. conceived the concept and designed the experiments. Z.X. conducted and analyzed the animal studies, flow cytometry, and imaging experiments. F.L. performed the single-cell RNA sequencing and bioinformatic analysis. Y.D., T.P., Z.G., and R.L.S. aided in the collection of clinical samples. Y.W. assisted in performing intra-iliac injections. X.H. and Y.G. assisted in performing intra-cardiac injections. J.L., I.L.B., and W.Z. contributed to maintaining mouse strains. W.L., L.Y., D.G.E., and S.A. provided assistance with animal work. H.L.C. aided in cell differentiation work. M.W.D., E.C., and other authors contributed to manuscript editing. X.H.-F.Z. supervised the research.

## SUPPLEMENTARY FIGURE LEGENDS

**Supplementary Figure 1.**
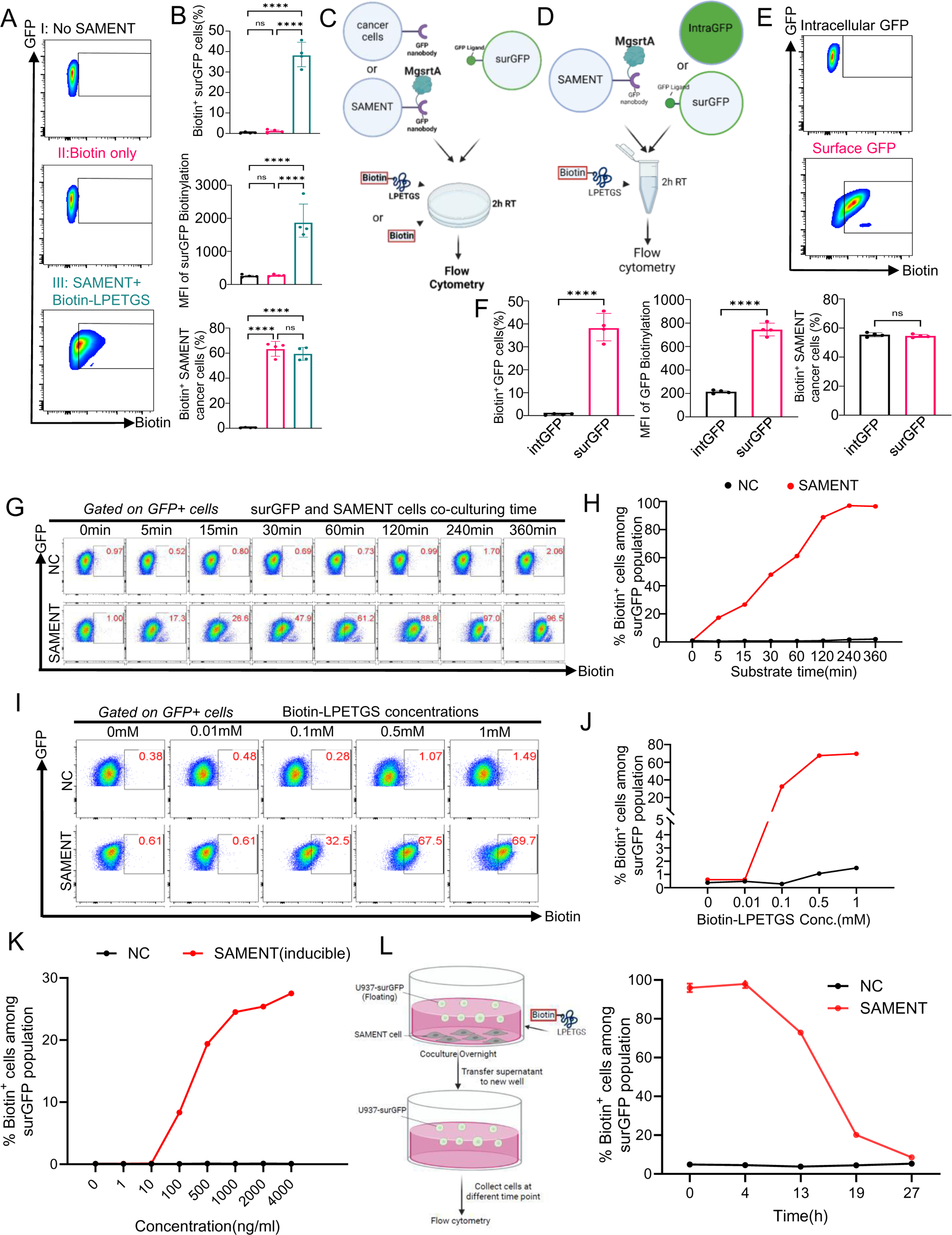
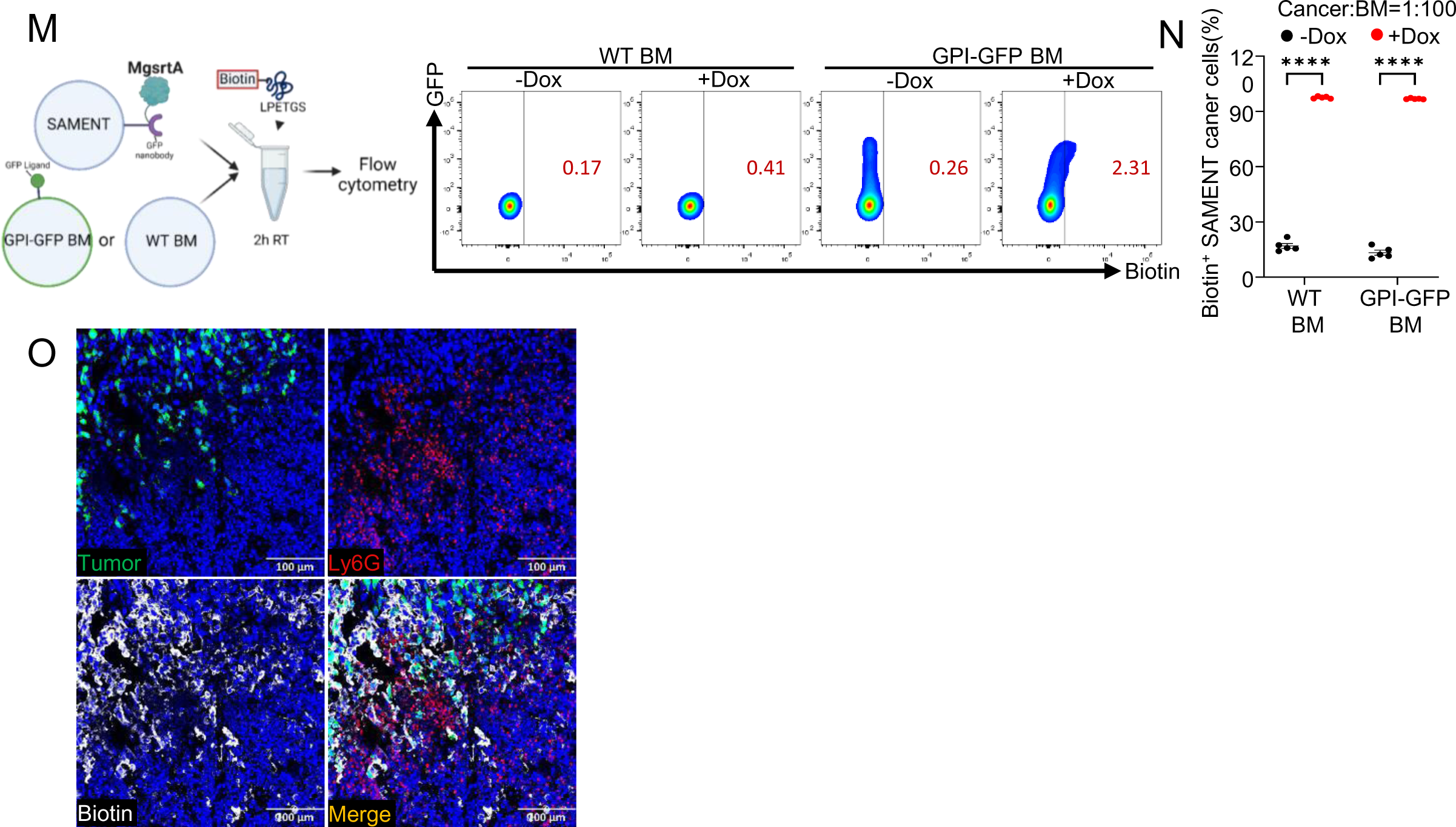
Validation and optimization of SAMENT *in vitro*, as related to Figure 1. **(A)** Representative FACS plots showing the biotinylation of surface-presented GFP (surGFP) receiver cells during co-culture: with normal cancer cells (top), with SAMENT cancer cells in the presence of biotin only (middle), or with SAMENT cancer cells in the presence of biotin-LPETGS (SAMENT substrate). **(B)** Percentage (top) and median fluorescence intensity (MFI) (middle) of biotin+ surGFP cells (gated as in A) among total surGFP cells. Percentage (Right) of biotin+ SAMENT cells among total SAMENT cell populations. This indicates that biotinylation of surGFP cells occurs only in the presence of both SAMENT and the suitable substrate, biotin-LPETGS. Four biological replicates per group were tested. **(C)** Schematic diagram illustrating the experimental setup (A-B) for labeling surGFP receiver cells during co-culture with SAMENT+ cancer cells or corresponding control cancer cells *in vitro*, followed by the addition of either biotin-LPETGS substrate or a biotin control group. **(D)** Schematic diagram illustrating the experimental setup(E-F) for labeling by SAMENT+ cancer cells when co-culturing with surGFP or intracellular GFP (IntraGFP) receiver cells *in vitro* **(E)** Representative FACS plots showing biotinylation of IntraGFP receiver cells (top) and surGFP receiver cells (bottom) through co-culturing with SAMENT^+^ cancer cells *in vitro*. **(F)** The percentage (left) and median fluorescence intensity (MFI, middle) of biotin+ surGFP cells (gated as in E) were notably higher than those of intraGFP cells. This observation suggests that surGFP cells demonstrate enhanced biotinylation by SAMENT, attributed to increased intercellular interaction when compared with intraGFP cells. Percentage (right) of biotin+ SAMENT cells among the total SAMENT cell populations, demonstrating similar substrate binding ability of SAMENT cells in both groups. Four biological replicates per group were tested. **(G)** Representative flow cytometric analysis showing biotinylation of surGFP cells by SAMENT with different co-culturing times. SAMENT cancer cells were co-cultured with surGFP cells at the ratio of 1:1 for the indicated time with the addition of biotin-LPETGS (500 uM) *in vitro*. **(H)** Quantification of data represented by (G). **(I)** Representative flow cytometric analysis depicting the biotinylation of surGFP cells by SAMENT with varying amounts of biotin-LPETGS substrate. SAMENT cancer cells were co-cultured with surGFP cells at a 1:1 ratio under the addition of the indicated concentrations of biotin-LPETGS, followed by a 2-hour incubation *in vitro*. **(J)** Quantification of data represented by (I). **(K)** Flow cytometric analysis shows the biotinylation of surGFP cells by tet-on inducible SAMENT using various concentrations of doxycycline for induction. Tet-on inducible SAMENT cancer cells or normal LLC1 cells (NC) were induced overnight with the indicated concentrations of doxycycline. Subsequently, surGFP cells were co-cultured with these cells at a 1:1 ratio, followed by the addition of 500uM biotin-LPETGS and a 2-hour incubation period *in vitro*. **(L)** Left: Schematic diagram illustrating the experimental setup (Right) for detecting the existence of biotinylation by SAMENT+ cancer cells on surGFP cells. The floating surGFP cell line U937, was co-cultured overnight with SAMENT cells under biotin-LPETGS treatment for complete labeling. The supernatant containing floating U937 cells was washed three times to remove unbound biotin-LPETGS. Subsequently, it was transferred to a new culture well and collected at different time points for flow cytometry analysis to measure the existing biotinylation levels. Right: The percentage of biotin+ cells among total U937-surGFP cells was measured at different time points after transferring U937-surGFP cells to a non-labeling environment. NC: normal LLC1 cells. SAMENT: SAMENT+ LLC1 cells. **(M)** Experimental setup (Left) and representative FACS plots (Right) of biotin labeling on GPI-GFP+ bone marrow cells isolated from GPI-GFP mice or WT bone marrow cells from WT mice, co-cultured *ex vivo* with SAMENT+ LLC1 cells. Flow cytometric analysis is referenced in Figure 1D. **(N)** Percentage of biotin+ SAMENT cells among the total SAMENT cell populations, demonstrating similar substrate binding ability of SAMENT cells in both groups. Flow cytometric analysis is referenced in Figure 1D. Five biological replicates per group were tested. **(O)** Fluorescence imaging captures SAMENT-mediated biotinylation of neutrophils (Ly6G^+^) in the LLC1 bone metastasis model. Data representative of two independent experiments. Data are presented as geometric mean ± geometric SD in B, F, N. Data are presented as mean ± SEM in L. P values were assessed by repeat measure one-way ANOVA followed by least significant difference (LSD) test in B; by unpaired t-test in F, N;

**Supplementary Figure 2.**
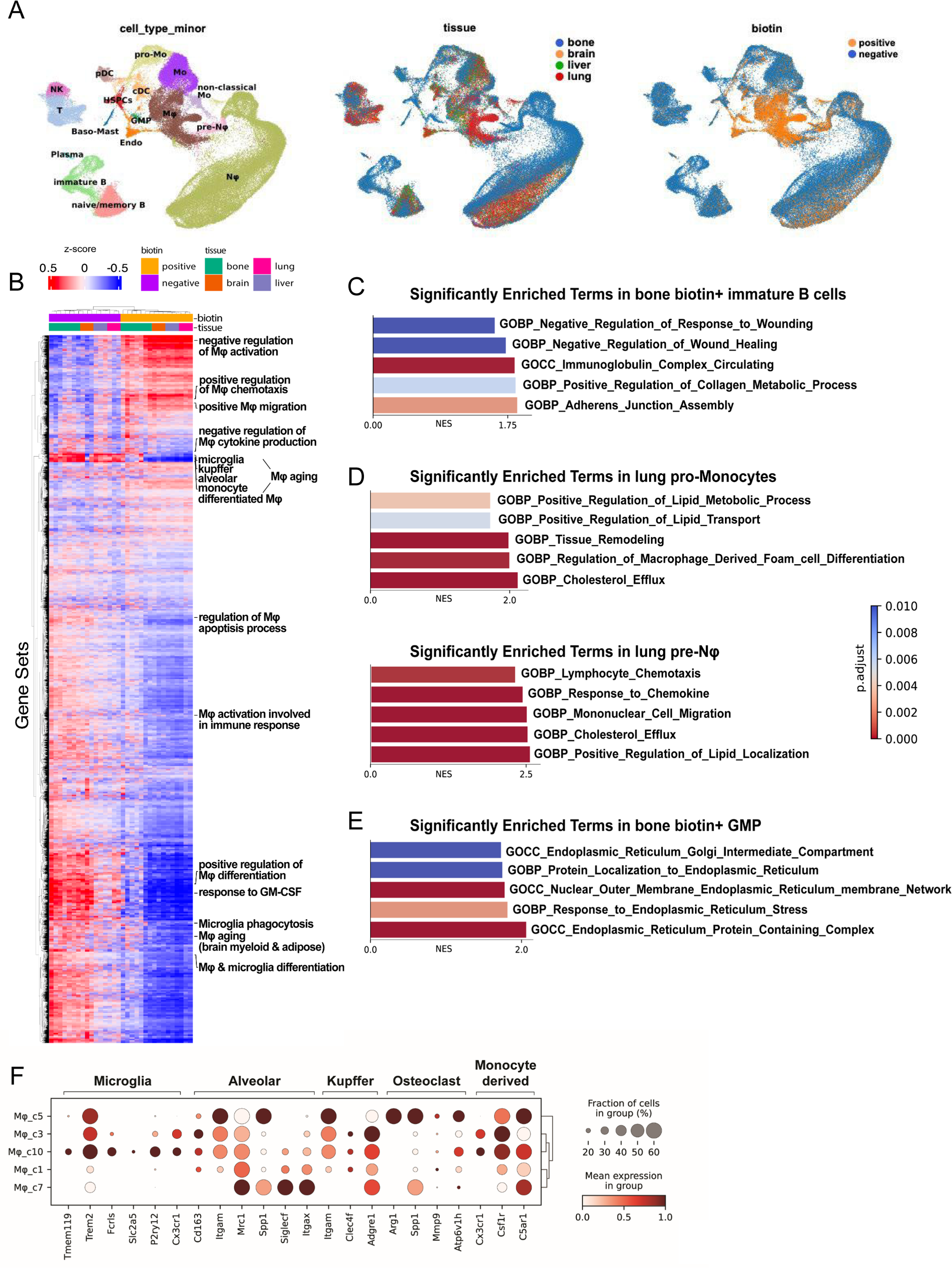
Additional examples of single-cell transcriptomic variations between biotin-positive and biotin-negative cell types, as related to Figure 2. **(A)** Cell types, tissue origin, and biotin groups in SAMENT single-cell dataset. **(B)** GSVA analysis comparing biotin-positive NK cells to biotin-negative NK cells highlights potential regulations exerted by NK cells on macrophages. Biotin-positive NK cells appear to modulate the tumor microenvironment by enhancing chemotaxis and inhibiting both aging and apoptosis in macrophages. This suggests that upon encountering tumors, reprogrammed NK cells indirectly contribute to chronic inflammation and promote the survival of tumor-associated macrophages. **(C)** Top enriched pathways in biotin-positive immature B cells in bones compared with other tissues. **(D)** Top enriched pathways in monocyte and neutrophil precursors in lungs compared with other tissues. **(E)** Top enriched pathways in biotin-positive granulocyte-macrophage progenitors (GMPs) in bone compared with biotin-negative populations. **(F)** Validation of published tissue-specific macrophage signatures was conducted in the SAMENT dataset. Reported Microglia signatures^50^ include Tmem119^+^, Trem2^+^, Fcrls^+^, Slc2a5^+^, P2ry12^+^, and Cx3cr1^+^. Alveolars signatures^51^ consist of Cd163^−^, Cd11b^−^, Cd206^int^, Spp1^+^, SiglecF^+^ and Cd11c^+^; Osteoclasts signatures^37^ are characterized by Arg1^+^, Spp1^+^, Mmp9^+^, and vacuolar [H+]-ATPase^+^; Kupffers cell signatures^52^ include CD11b^low^, F4/80^high^, and Clec4F^+^; Monocyte-derived macrophage signatures^52,53^ encompass CD11b^+^, F4/80^int^, Ly6C^+^ and CSF1R^+^. In our dataset, Mφ_c10 corresponds to Microglia, Mφ_c7 corresponds Alveolar macrophages, and Mφ_c5 corresponds to osteoclasts. Monocyte-derived macrophages are distributed across all macrophage subclusters, while we did not identify specific clusters for Kupffer cells. These findings support the observations depicted in Figure 2I.

**Supplementary Figure 3.**
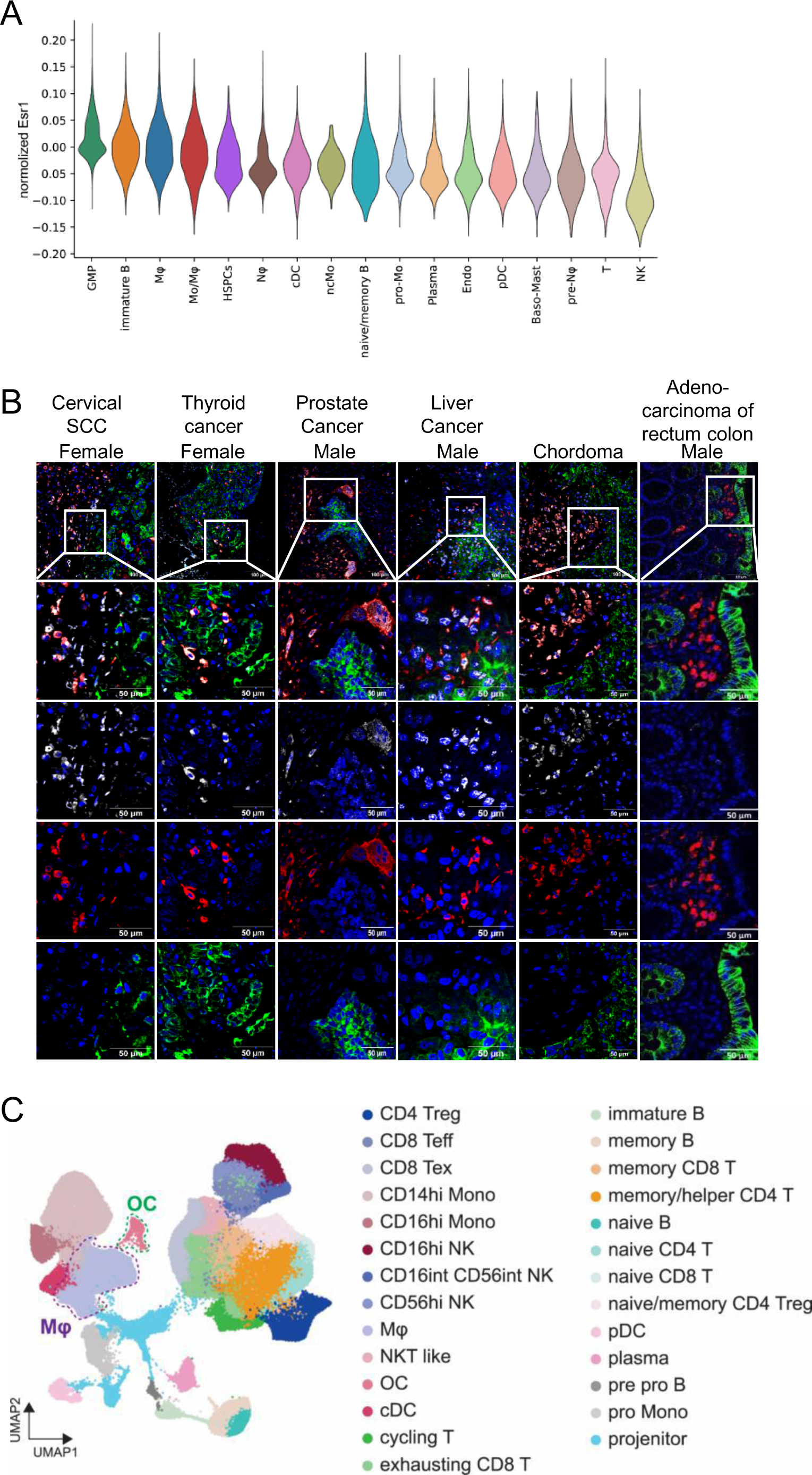
Ranked *Esr1* expression across cell types in biotin-positive bone niche cells, as related to Figure 3. **(A)** The normalized and scaled *Esr1* gene expression across all cell types in the biotin-positive niche is presented. The violin plot is ranked based on the median *Esr1* expression levels. Macrophages exhibit high levels of *Esr1* expression. **(B)** Immunofluorescence staining of cytokeratin 8/19-positive cancer cells in diverse bone metastases tumors demonstrates co-localization of ERα and CD68 (a human macrophage marker). Notably, in cases of colon tumor metastasis, CD68+ macrophages exhibit very low expression of ERα. Data representative of at least two independent patient samples staining. **(C)** UMAP projections depict the human bone metastasis single-cell RNA sequencing dataset, with cells color-coded based on their lineage annotation.

**Supplementary Figure 4.**
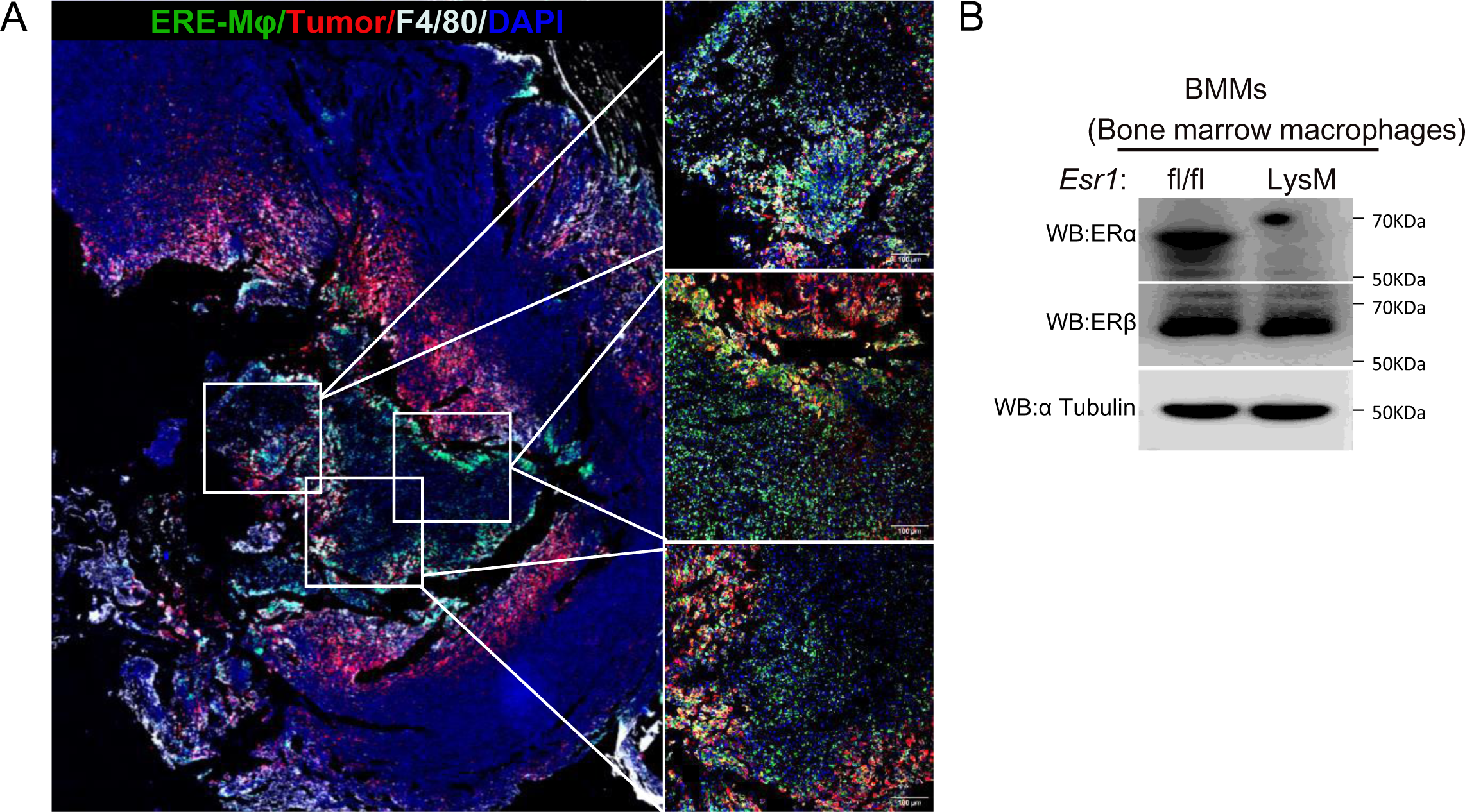
Interaction with metastatic cancer cells induces the expression and activity of ERα, as related to Figure 4. **(A)** Bone marrow-derived macrophages were infected with the ERE-Luc-T2A-GFP reporter virus and subsequently transplanted into macrophage-depleted WT mice via retro-orbital injection. Bones were collected four days after transplantation. Immunofluorescence staining revealed an increased presence of ERE-Luc-T2A-GFP-infected macrophages in close proximity to tumor cells. **(B)** A representative Western blot depicting the expression of ERα, ERβ and the internal control, α-tubulin, in macrophages derived from *Esr1*^fl/fl^ and LysM-cre *Esr1*^fl/fl^ bone marrow.

**Supplementary Figure 5.**
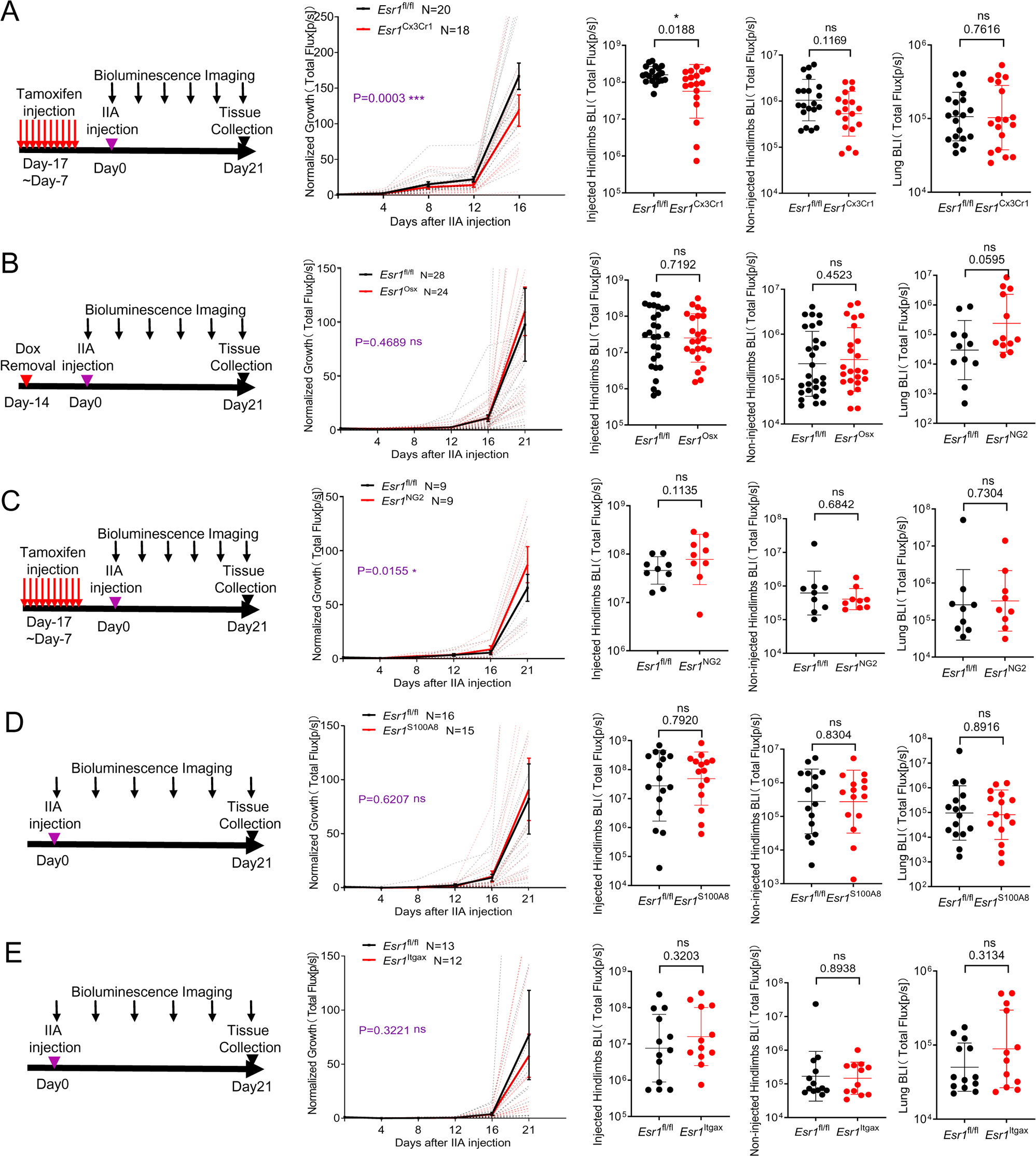
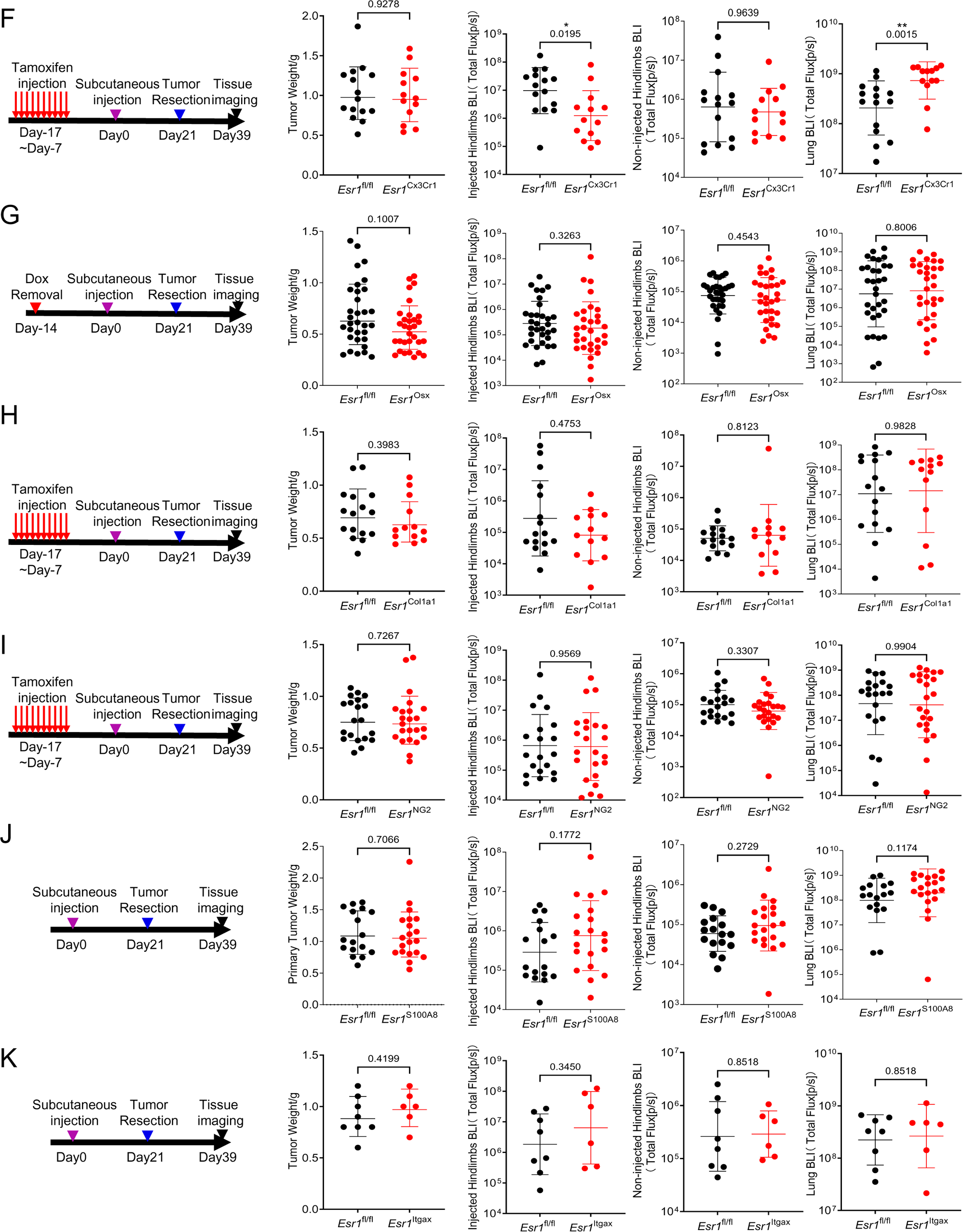
Depleting ERα in different bone stromal cells or myeloid lineage cells leads to variable impacts in experimental metastasis assays and spontaneous metastasis assays, as related to Figure 5. **(A)** The schematic diagram (left), normalized tumor growth curves(middle), and quantified *ex vivo* bioluminescence imaging intensities in lungs or hindlimb bones(right three) of female Cx3Cr1-CreERT2 *Esr1*^fl/fl^ mice (n=18 mice) and corresponding control mice (n=20 mice) are presented. Cx3Cr1 serves as macrophage lineage marker. **(B)** The schematic diagram (left), normalized tumor growth curves(middle), and quantified *ex vivo* bioluminescence imaging intensities in lungs or hindlimb bones(right three) of female Osx-tTA-Cre *Esr1*^fl/fl^ mice (n=22 mice) and corresponding control mice (n=25 mice) are presented. Osx serves as osteoprogenitor cell marker. **(C)** The schematic diagram (left), normalized tumor growth curves(middle), and quantified *ex vivo* bioluminescence imaging intensities in lungs or hindlimb bones(right three) of female NG2-CreERT2 *Esr1*^fl/fl^ mice (n=9 mice) and corresponding control mice (n=9 mice) are presented. NG2 serves as a marker for mesenchymal stem cells and pericytes. **(D)** The schematic diagram (left), normalized tumor growth curves(middle), and quantified *ex vivo* bioluminescence imaging intensities in lungs or hindlimb bones(right three) of female S100A8-Cre *Esr1*^fl/fl^ mice (n=15 mice) and corresponding control mice (n=16 mice) are presented. S100A8 serves as neutrophil cell marker. **(E)** The schematic diagram (left), normalized tumor growth curves(middle), and quantified *ex vivo* bioluminescence imaging intensities in lungs or hindlimb bones(right three) of female Itgax-Cre *Esr1*^fl/fl^ mice (n=12 mice) and corresponding control mice (n=13 mice) are presented. Itgax serves as dendritic cell marker. **(F)** The schematic diagram(left), tumor weight(middle), and spontaneous metastasis to lungs or hindlimb bones(right three) in female Cx3Cr1-CreERT2 *Esr1*^fl/fl^ (n=13 mice) and corresponding control mice (n=15 mice) are presented. Cx3Cr1 serves as macrophage lineage marker. **(G)** The schematic diagram(left), tumor weight(middle), and spontaneous metastasis to lungs or hindlimb bones(right three) in female Osx-tTA-Cre *Esr1*^fl/fl^ (n=31 mice) and corresponding control mice (n=32 mice) are presented. Osx serves as osteoprogenitor cell marker. **(H)** The schematic diagram(left), tumor weight(middle), and spontaneous metastasis to lungs or hindlimb bones(right three) in female Col1a1-CreERT2 *Esr1*^fl/fl^(n=13 mice) and corresponding control mice (n=16 mice) are presented. Col1a1 serves as a marker for differentiated osteoblasts. **(I)** The schematic diagram(left), tumor weight(middle), and spontaneous metastasis to lungs or hindlimb bones(right three) in female NG2-CreERT2 *Esr1*^fl/fl^(n=23 mice) and corresponding control mice (n=20 mice) are presented. NG2 serves as a marker for mesenchymal stem cells and pericytes. **(J)** The schematic diagram(left), tumor weight(middle), and spontaneous metastasis to lungs or hindlimb bones(right three) in female S100A8-Cre *Esr1*^fl/fl^(n=20 mice) and corresponding control mice (n=17 mice) are presented. S100A8 serves as neutrophil cell marker. **(K)** The schematic diagram(left), tumor weight(middle), and spontaneous metastasis to lungs or hindlimb bones(right three) in female Itgax-Cre *Esr1*^fl/fl^(n=6 mice) and corresponding control mice (n=8 mice) are presented. Itgax serves as dendritic cell marker. Each dotted curve represents an individual animal, whereas the highlighted curve shows the mean growth for each group in A, B, C, D(middle). Data are presented as mean ± SEM for all tumor growth curves; geometric mean ± geometric SD for all scatter plot. P values were assessed by Fisher least significant difference test post repeat measure two-way ANOVA test for all tumor growth curves; by Mann–Whitney test for all scatter plots.

**Supplementary Figure 6.**
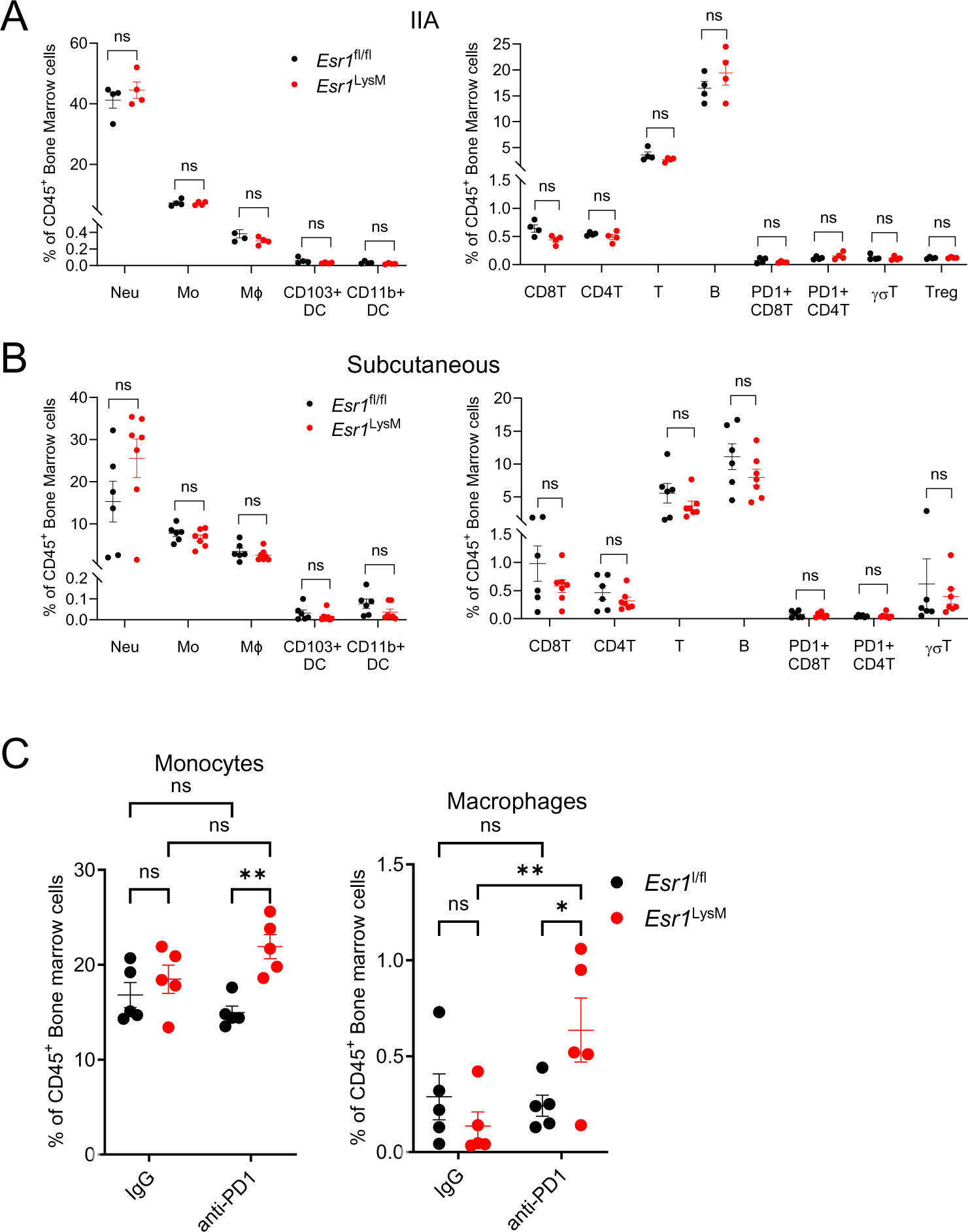
The impact of *Esr1* knockout on immune cell frequency, as related to Figure 6. **(A)** Quantification of major myeloid (left) and lymphoid (right) cell populations in the hindlimb bones of female LysM-Cre *Esr1*^fl/fl^ mice (n=4 mice) and their corresponding control mice (n=4 mice) after IIA injection with LLC1 cells. **(B)** Quantification of primary myeloid (left) and lymphoid (right) cell populations in the hindlimb bones of female LysM-Cre *Esr1*^fl/fl^ mice (n=7 mice) and their corresponding controls (n=6 mice) with spontaneous bone metastasis in LLC1 models. **(C)** Monocyte and macrophage quantification in the hindlimb bones of mice injected with LLC1 cells, including LysM-Cre *Esr1*^fl/fl^ mice (n=5 mice) and their control *Esr1*^fl/fl^counterparts (n=5 mice), with or without anti-PD1 treatment after IIA injection. Data are presented as mean ± SEM in A, B, C; P values were assessed by Multiple Mann–Whitney test in A, B; by Fisher least significant difference test post repeat measure two-way ANOVA test in C.

## STAR METHODS

### KEY RESOURCES TABLE

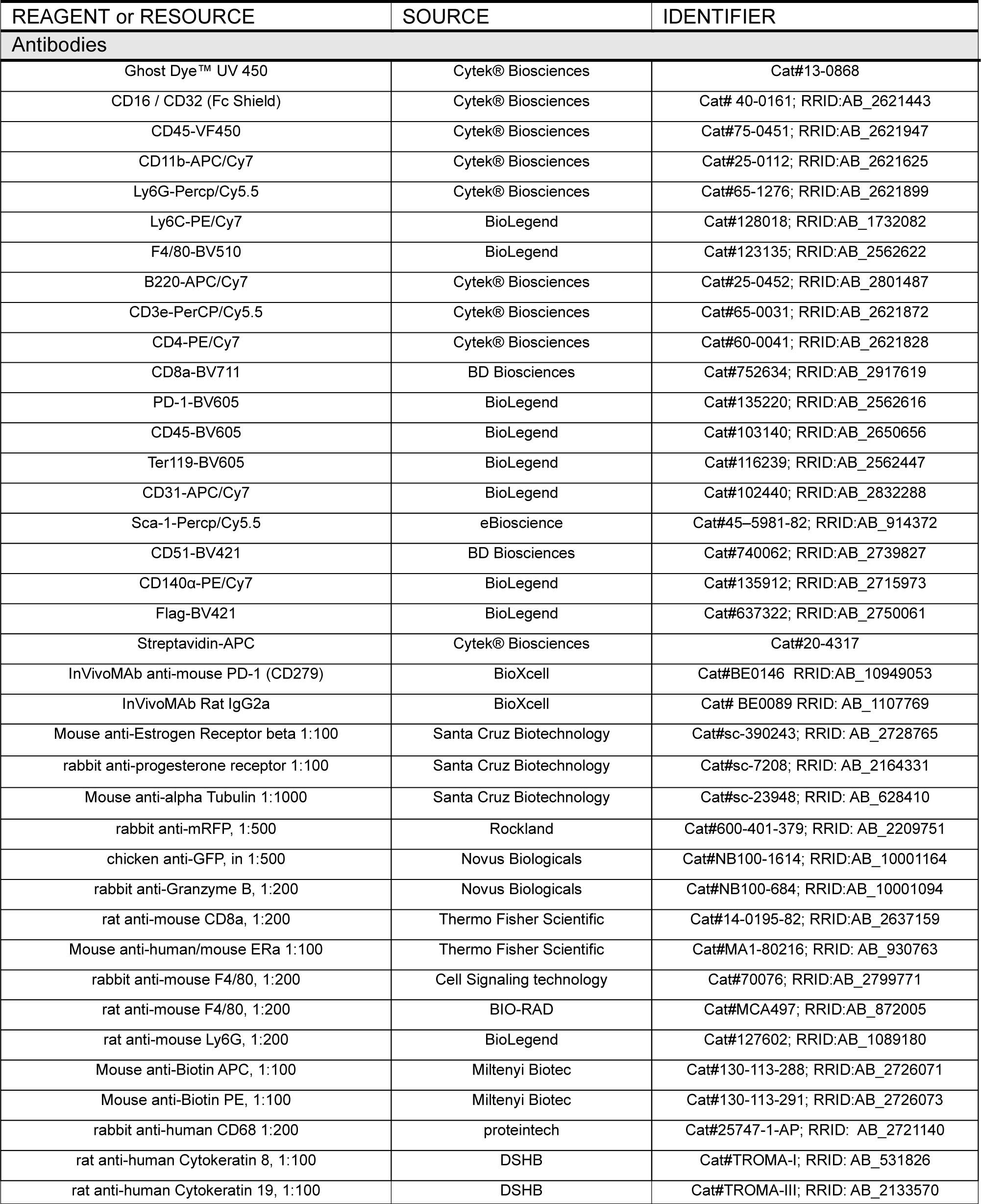
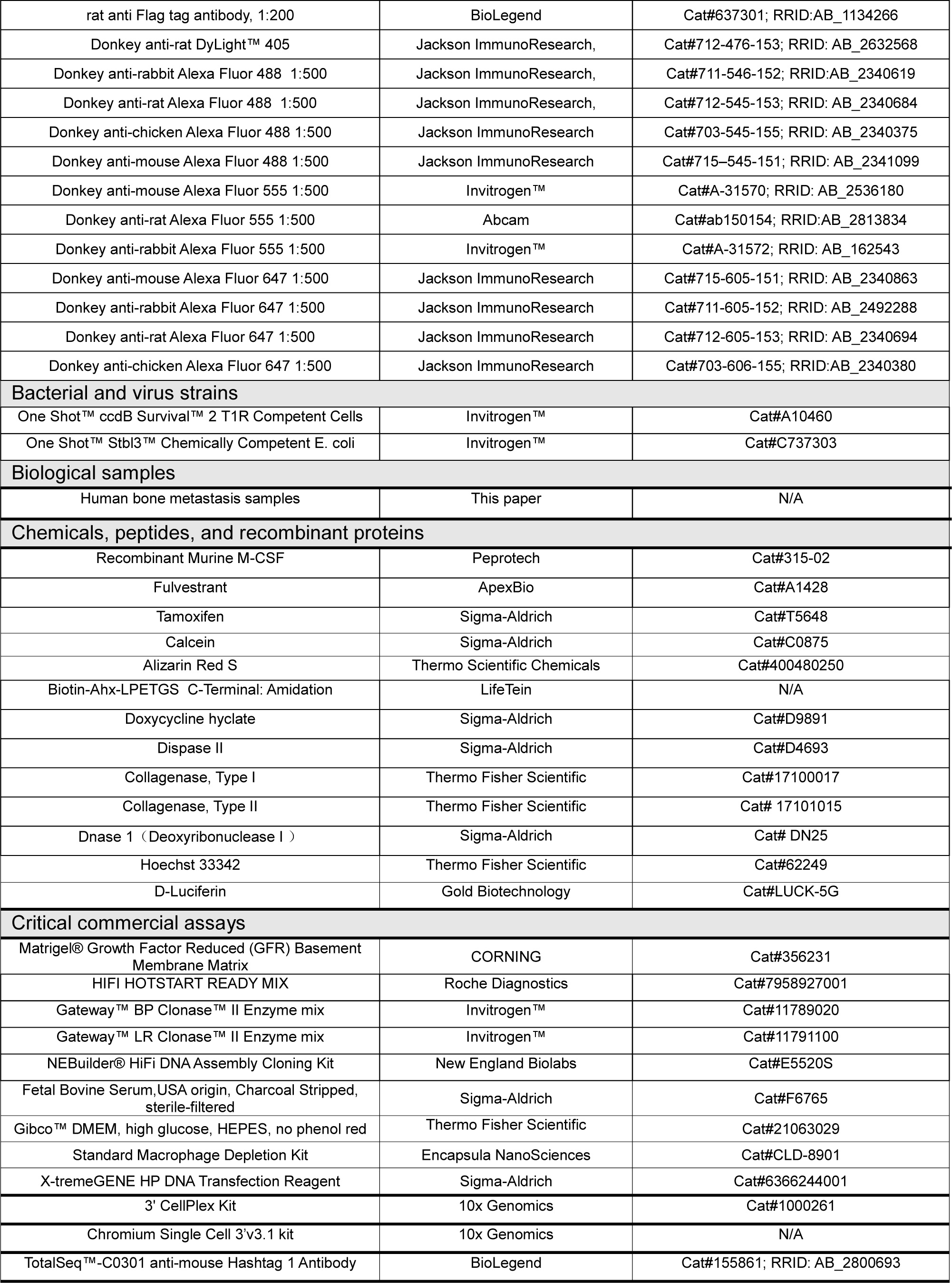
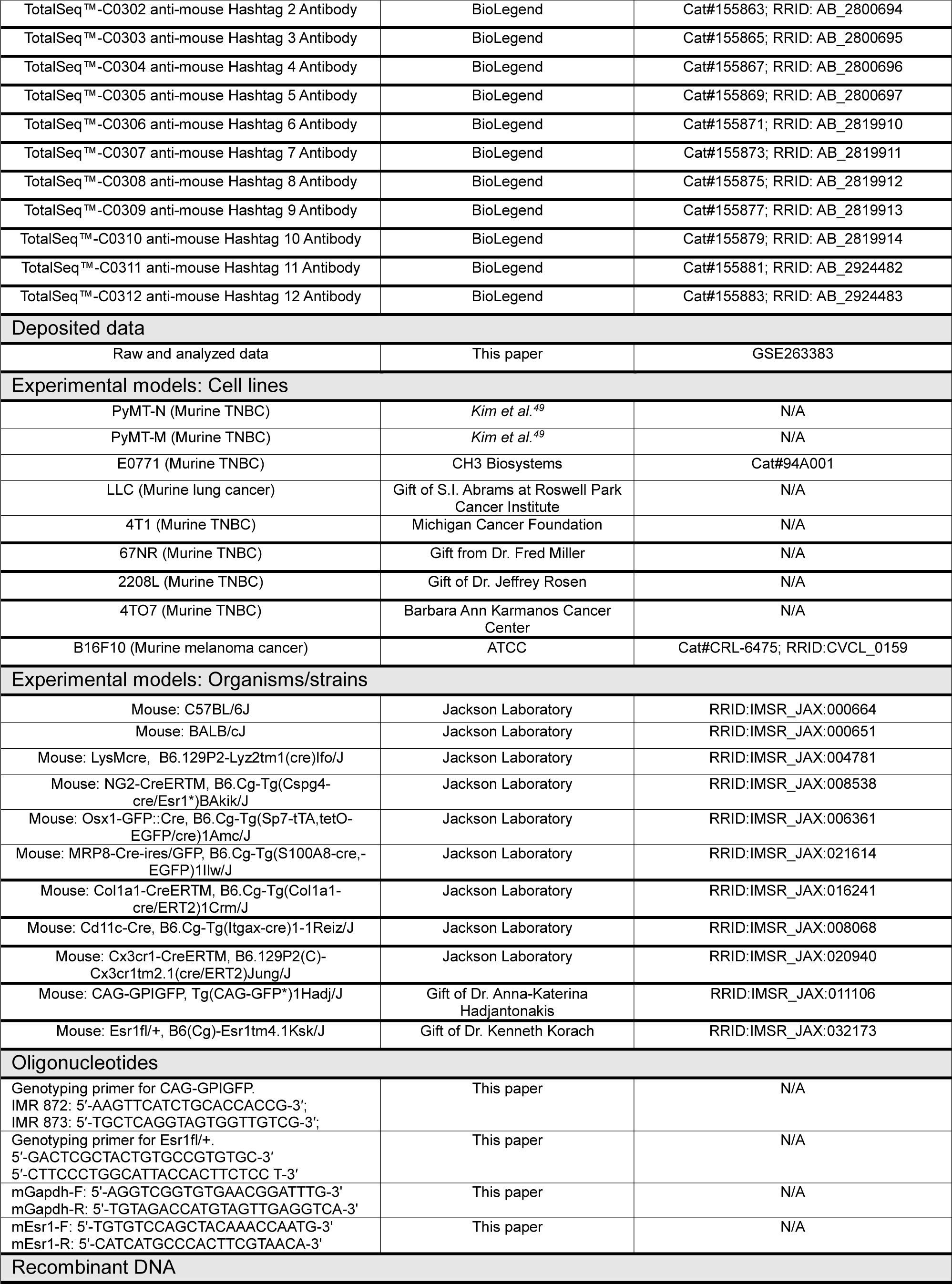
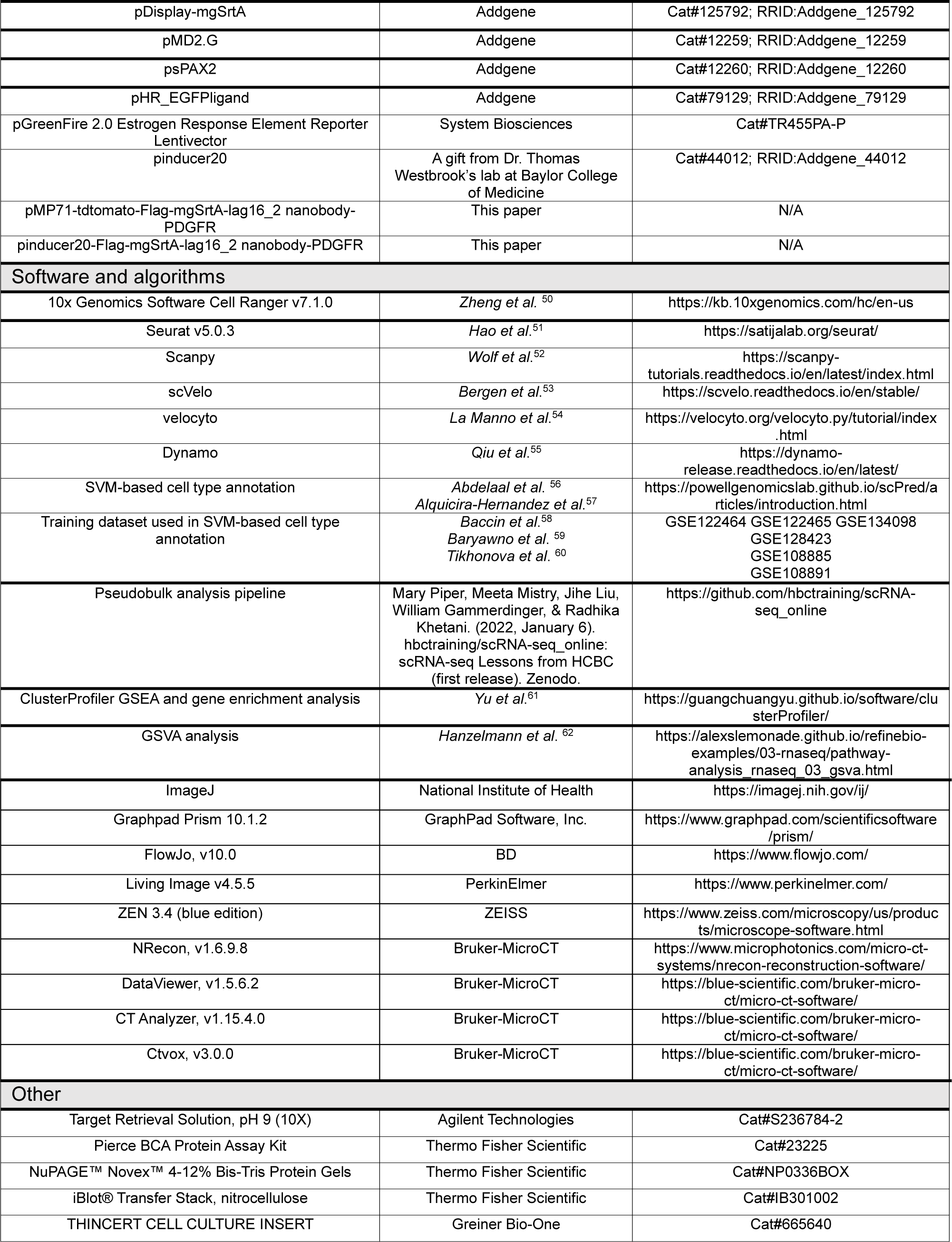

### RESOURCE AVAILABILITY

#### Lead Contact

Further information and requests for resources and reagents should be directed to and will be fulfilled by the Lead Contact, Xiang H.-F. Zhang.

#### Materials Availability

All unique/stable reagents generated in this study are available from the Lead Contact with a completed Materials Transfer Agreement.

#### Data and Code Availability

Raw and analyzed scRNA-seq data has been deposited in the NCBI GEO database (accession number: GSE263383).

#### Supplementary table

This table contains the DNA sequences of the plasmids used in the study.

### EXPERIMENTAL MODEL AND SUBJECT DETAILS

#### Mouse Model

The *in vivo* procedures and use of animal models were conducted following protocol AN-5734 approved by the Baylor College of Medicine Institutional Animal Care and Use Committee. The mouse strains utilized, including C57BL/6J (RRID:IMSR_JAX:000664), BALB/cJ (RRID: IMSR_JAX:000651), LysMcre (RRID:IMSR_JAX:004781), NG2-CreERTM (RRID:IMSR_JAX:008538), Osx1-GFP::Cre (RRID:IMSR_JAX:006361), S100A8-Cre-ires/GFP (RRID:IMSR_JAX:021614), Col1a1-CreERTM (RRID:IMSR_JAX:016241), Cd11c-Cre (RRID:IMSR_JAX:008068), and Cx3cr1-CreERTM (RRID:IMSR_JAX:020940), were obtained from The Jackson Laboratory. Mice genotyping was performed according to the JAX genotyping protocol. The CAG-GPIGFP strain (C57BL/6J, RRID:IMSR_JAX:011106) was generously provided by Dr. Anna-Katerina Hadjantonakis^20^ (Memorial Sloan-Kettering Cancer Center), and *Esr1*^fl/+^ mice(RRID:IMSR_JAX:032173) were provided by Dr. Kenneth Korach^69^ (National Institute of Environmental Health Sciences). Detailed information on the primers used for genotyping is available in the **key resource table**. Conditional *Esr1* knockout strains were generated through crosses between male Cre+ transgenic mice and female *Esr1*^fl/fl^ mice over two generations. Additionally, B6.LysMcre, B6. *Esr1*^fl/fl^, and B6.CAG-GFPGPI mice underwent over 10 generations of backcrossing to BALB/cJ in our lab, followed by further breeding to establish the BALB/cJ-LysM-Cre+/−; *Esr1*^fl/fl^ strain.

#### Human Bone Metastasis Samples

The collection of human bone metastasis samples adhered to the Declaration of Helsinki guidelines and received approval from the Institutional Review Boards at Baylor College of Medicine (H-49396), The University of Texas MD Anderson Cancer Center (PA15-0225), and the University of Texas Medical Branch (H-46675). Written informed consent was obtained from all patients undergoing orthopedic surgery, authorizing the research use of their samples.

#### Cell lines

Murine triple-negative breast cancer (TNBC) lines, including PyMT-N (B6), PyMT-M (B6), 4T1 (BALB/c), 4TO7 (BALB/c), 67NR (BALB/c), and 2208L (BALB/c), along with the Mouse Lewis lung carcinoma cell line: LLC (B6) and Mouse melanoma cell line: B16F10 (B6), were cultured in DMEM/high glucose medium (Cytiva, Cat#SH30022.01) supplemented with 10% FBS (Thermo Fisher Scientific, Cat#A5256701) and antibiotics(fisher scientific, Cat#MT30004CI). Additionally, 67NR (BALB/c) cells received NEAA supplementation. The TNBC line E0771 (B6) was cultured in RPMI-1640 medium (Cytiva, Cat#SH30027.01) supplemented with 10% FBS, 1% HEPES (Cytiva, Cat#SH30237.01), and antibiotics.

### METHOD DETAILS

#### Plasmids, Virus Production, and Infection

The SAMENT sequence was constructed by combining various elements, including an N-terminal Mouse Ig Kappa signal peptide, Flag-tag, mgSrtA^12^, Lag16_2 GFP nanobody^19^ and the PDGFRβ transmembrane domain. This SAMENT sequence was then inserted into the pMP71 vector for continuous expression or into the pinducer20 vector^70^ for inducible expression. The pMP71 construct was assembled using NEBuilder DNA assembly (New England Biolabs, E5520S), while the Pinducer20 constructs were generated through Gateway cloning. Details of all cloned constructs are provided in Supplementary Table 1.

To produce lentivirus or retrovirus, the plasmids were transfected into HEK293T cells along with packaging plasmids using X-tremeGENE HP DNA Transfection Reagent (Sigma-Aldrich, 6366244001). After 48 hours, the supernatant containing the viral particles was collected and filtered using a 0.45 μm filter. Cells were then exposed to the lentivirus or retrovirus along with 10 μg/ml polybrene (EMD Millipore,TR-1003-G) to facilitate infection. Postive cells were sorted to establish the desired cell lines.

#### SAMENT *In vitro* labeling

SAMENT- and surGFP-expressing cells were cultured individually in DMEM medium. Inducible SAMENT cells were pre-treated with 1 μg/ml Doxycycline for 12 hours before collection. Following detachment with Trypsin, the cells were washed and resuspended in 100 μL PBS. Subsequently, these two cell populations were mixed in a 1.5-ml microcentrifuge tube, and biotin-LPETGS was added at a final concentration of 500 μM or the specified concentration. After a 2-hour co-culture or the specified duration at room temperature, cells were washed three times with PBS to remove excess biotin probes. The cell mixture was then incubated with Streptavidin-APC (Cytek® Biosciences, 20-4317) and Flag-VF450 (BioLegend,637322) on ice for 15 minutes before undergoing PBS washes for subsequent FACS analysis.

#### SAMENT *In vivo* labeling

Mice receiving IIA or IC injections were administered 2 mg of doxycycline via intraperitoneal injection for two consecutive days prior to euthanization. Additionally, a solution of 2 mM biotin-LPETGS (LifeTein, dissolved in PBS at a concentration of 20 mM) was injected via retroorbital injection 12 hours before and again 2 hours before euthanization.

#### Induction of Cre-Mediated Recombination

If not stated otherwise, female mice were used for all *in vivo* experiments, with age-matched Cre−/− littermates serving as controls. To activate Cre-ERT2, female mice aged 5 weeks received daily injections of 1 mg tamoxifen (Sigma-Aldrich, T5648) dissolved in a solution comprising 5% ethanol and 95% corn oil (Sigma-Aldrich, C8267). This regimen was administered for 10 consecutive days. In the case of Osx1-GFP::Cre mice, expression of the EGFP/Cre fusion protein was suppressed by adding doxycycline to water bottles at a final concentration of 0.2mg/ml, which was changed weekly until 2 weeks before the experiment.

#### MicroCT Analysis of Bone Samples

The femur bones were initially prepared by removing the skin and then fixed in 70% ethanol before undergoing scanning using a Bruker Skyscan 1272 scanner (Bruker-MicroCT). The X-ray energy was set at 50 kV and 200 μA, with images captured at a pixel size of 6.6 μm. A rotation angle of 0.1° was employed for a complete 360° rotation during scanning. Trabecular bone analysis focused on a 2.0 mm region of the distal metaphysis, beginning 0.7 mm from the proximal end of the distal femoral growth plate, with a threshold range set at 75–255 permille. For cortical bone analysis, a 0.5 mm region of the femoral cortical bone, starting 3.7 mm from the proximal end of the distal femoral growth plate, was assessed using a threshold range of 125–255 permille. Cross-sectional images were generated using NRecon (Bruker-MicroCT, v1.6.9.8) and DataViewer (Bruker-MicroCT, v1.5.6.2), while bone volumes were analyzed via CT Analyzer (Bruker-MicroCT, v1.15.4.0). Reconstruction of three-dimensional bone images was performed using CTvox (Bruker-MicroCT, v3.0.0).

#### Bone Formation Rate Measurement by Calcein-alizarin red S labeling

Ten-week-old mice were intraperitoneally injected with Calcein (Sigma-Aldrich, C0875) at 20 mg/kg on day 0 and with Alizarin Red S (Thermo Scientific Chemicals, 400480250) at 40 mg/kg on day 4. Upon euthanasia on day 7, hindlimb bones were collected, fixed overnight, and snap-frozen in OCT. Non-decalcified bone sections were obtained using the CryoJane tape-transfer system on a Leica CM3050S Cryostat equipped with Low-Profile Disposable Blades DB80LX (Leica Biosystems, 14035843496). These sections were then mounted with Prolong Gold Antifade Mountant with DAPI (Invitrogen, P36935). Imaging of the distance between Calcein and Alizarin Red S labeling was conducted using an LSM880 confocal microscope, followed by analysis using ImageJ.

#### Intra-Iliac Artery (IIA) injection and Intra-Cardiac (IC) injection

Intra-iliac artery injections were performed following established protocols^21^. After anesthetizing the animals and sterilizing the surgical site, a small incision (7-8 mm) was made between the fourth and fifth nipples to access the iliac vessels. Cancer cells, suspended in 100μL of PBS, were then slowly injected into the iliac artery using a 31-G insulin syringe (Becton Dickinson, 328418). Gentle pressure was applied with cotton tips to manage any bleeding before closing the wound. For intra-cardiac injection, cancer cells suspended in 100μL of PBS were directly administered into the left ventricle of anesthetized animals using a 26-gauge syringe (Becton Dickinson, 309625). The needle was carefully inserted into the heart, followed by gradual withdrawal, and pressure was applied to minimize bleeding.

#### Implantation of Tumors in the Mammary Fat Pad or Subcutaneously and Surgical Tumor Removal

The LLC1 cells were suspended in PBS and combined with growth factor-reduced matrigel (Corning, Cat#356231) at a 1:1 ratio. An equivalent volume of growth factor–reduced matrigel was mixed with the cells and administered via subcutaneous injection into the skin near the left hindlimb. Tumor removal surgery was conducted approximately 3 weeks after implantation to completely excise the primary tumors. Most animals survived for about 2-3 weeks after the removal of primary tumors and were subsequently dissected and examined between days 37 and 40 for lung and bone metastasis.

#### Administration of Fulvestrant and Immune Checkpoint Blockade (ICB) treatment

For Fulvestrant treatment, the selective estrogen receptor degrader (SERD) fulvestrant (ApexBio, #A1428) was dissolved in a solution consisting of 5% DMSO and 95% corn oil (Sigma-Aldrich, C8267). Mice were subcutaneously administered with a dose of 250 mg/kg per mouse starting from day 3 after IIA injection. This treatment was given once a week for a total duration of 3 weeks. For ICB treatment, mice received intraperitoneal injections of either anti-PD1 (200 mg per mouse,BioXcell,BE0146) or anti-IgG2a (200 mg per mouse,BioXcell,BE0089) every three days, commencing from day 1 post IIA injection. This dosing regimen was repeated for a total of six doses.

#### Bioluminescence Imaging and Tissue Collection

For *in vivo* bioluminescence imaging, mice in the IIA model were scanned using mode E of the IVIS Lumina II system (PerkinElmer) at predetermined intervals. Prior to imaging, the fur on the posterior aspect of the injected hindlimb was carefully removed to enhance the penetration of bioluminescence. Subsequently, animals received a retro-orbital injection of 100 μL of 15 mg/mL D-luciferin (Gold Biotechnology, LUCK-5G).

For ex vivo bioluminescence imaging, mice from either the IIA model or the spontaneous metastasis model were used. Dissected tissues were promptly scanned in D mode, with exposure time adjusted to avoid signal saturation. To quantify the total bioluminescence intensity of specific tissues, a standardized region of interest (ROI) was consistently applied to all animals or dissected tissues, and the total flux (photons/second) was quantified.

The dissected bones or soft tissues were immediately snap-frozen or placed in 10% neutral buffered formalin overnight. After fixation, the bone tissues were washed in PBS to remove residual formalin, followed by decalcification in a 14% pH 7.4 EDTA solution for 7 days. Subsequently, the tissues were cryopreserved in a 30% sucrose-PBS solution and then embedded in OCT (Tissue-Tek).

#### Single-cell Resuspension Preparation

The bone marrow cells were extracted from the tibia and femur bones using a 26-G syringe containing FACS buffer (PBS with 2% FBS, antibiotics, and 2mM EDTA). The remaining bones were then crushed thoroughly with a mortar and pestle. The fragmented bones underwent digestion in DMEM containing 1 mg/mL collagenase I (Thermo Fisher Scientific,17100017), 1 mg/mL collagenase II (Thermo Fisher Scientific,17101015), 3 mg/mL Dispase II (Sigma-Aldrich,D4693), 1 mg/mL BSA, and 0.1 mg/mL DNase I (Sigma-Aldrich, DN25) at 37°C for 1 hour on a shaker. After digestion, the bones were washed with FACS buffer. The cells released from the enzyme digestion and FACS washing were combined with bone marrow cells, filtered through a 70 mm strainer, lysed in red blood cell (RBC) lysis buffer (Cytek® Biosciences,TNB-4300-L100) and centrifuged for subsequent analysis.

For soft tissues, they were also crushed thoroughly with a mortar and pestle, followed by digestion in the same DMEM solution. After digestion, the tissues were washed with FACS buffer, filtered through a 70 mm strainer, lysed in red blood cell (RBC) lysis buffer (Cytek® Biosciences,TNB-4300-L100), and centrifuged for subsequent analysis.

#### Flow Cytometry

For the pooled bone marrow cells, staining was initiated with Ghost Dye UV450 (Cytek® Biosciences, 13–0868) in 1 mL PBS for 30 minutes. Following centrifugation, the cells were resuspended in 1 mL FACS buffer. Subsequently, samples were divided for staining with various panels and pre-treated with anti-CD16/32 antibody (Cytek® Biosciences, 40-0161) for 10 minutes on ice to block non-specific binding. This was followed by incubation with fluorescent dye–conjugated primary antibodies on ice for 15 minutes. The antibodies used in this study included: Immune cell panel, CD45-VF450 (Cytek® Biosciences,75–0451), CD11b-APC/Cy7 (Cytek® Biosciences, 25–0112), Ly6G-Percp/Cy5.5 (Cytek® Biosciences, 65–1276), Ly6C-PE/Cy7 (BioLegend,128018), F4/80-BV510 (BioLegend,123135), B220-APC/Cy7 (Cytek® Biosciences, 25-0452), CD3e-Percp/Cy5.5 (Cytek® Biosciences,65-0031), CD4-PE/Cy7 (Cytek® Biosciences,60-0041), CD8a-BV711 (BD Biosciences,752634), PD-1-BV605 (BioLegend,135219), Biotin-APC (Miltenyi Biotec,130-113-288) or Biotin-PE(Miltenyi Biotec,130-113-291); Stromal cell panel, CD45-BV605 (BioLegend,103140), Ter119-BV605 (BioLegend,116239), CD31-APC/Cy7 (BioLegend,102440), Sca-1-Percp/Cy5.5 (eBioscience,45–5981-82), CD51-BV421 (BD Biosciences, 740062), CD140α-PE/Cy7 (BioLegend, 135912), Biotin-APC (Miltenyi Biotec,130-113-288) or Biotin-PE(Miltenyi Biotec,130-113-291)

Flow cytometry analysis was conducted using a BD LSRFortessa flow cytometer and data were further analyzed with FlowJo software. The gating strategy for each population was as follows: Monocytes: DAPI−CD45+CD11b+Ly6C+Ly6G−; Neutrophils: DAPI−CD45+CD11b+Ly6C^mid-low^Ly6G+; Macrophages: DAPI−CD45+CD11b+Ly6C−Ly6G−F4/80+; B cells: DAPI−CD45+B220+; T cells: DAPI−CD45+CD3e+; CD4 T cells: DAPI−CD45+CD3e+CD4+; CD8a T cells: DAPI−CD45+CD3e+CD8a+; Endothelial cells: CD45-Ter119-CD31+; Osteoprogenitors: CD45-Ter119-CD31-CD51+CD140a+; Mature osteoblasts: CD45-Ter119-CD31-CD51+Sca1-; MSCs: CD45 Ter119-CD31-CD140a+;

#### Single-cell RNA Library Construction and Sequencing

Mice received SAMENT+ LLC1 cell injections either via intracardiac or intra-iliac routes. Following this, the previously described SAMENT *in vivo* labeling procedure was conducted before euthanization. Afterwards, tumor tissues from mice, encompassing lungs, livers, brains, and hindlimbs, were collected, and single-cell suspensions were prepared as previously outlined. Prior to sorting, debris removal was undertaken to minimize excessive cell debris contamination. Subsequently, cells were stained with Ghost Dye UV450 and Biotin flow antibody. The tissue cells were then sorted into Biotin+ and Biotin-populations using Arial. To optimize cost-effectiveness, we implemented a cell hashing strategy during sample processing and sequencing. Two distinct hashing methods were employed for two separate batches of single-cell experiments: CellPlex by 10X Genomics and TotalSeq C by BioLegend. Both approaches followed the Chromium Next GEM Single Cell 3ʹ v3.1 (10x Genomics): Cell Multiplexing protocol (CG000383) and were carried out in collaboration with the Single Cell Genomics Core at BCM. The resulting libraries, containing barcoded single-cell transcriptomes, were subjected to sequencing to a depth of 600 million reads using the Novoseq 6000 system (Illumina) at the Genomic and RNA Profiling Core (GARP) and Novogene. Data processing was performed using the CASAVA 1.8.1 pipeline (Illumina). Sequence reads were converted into FASTQ files, cell multiplexing oligo (CMO), antibody barcoding matrices, and unique molecular identifier (UMI) read counts using the CellRanger software from 10X Genomics.

#### Immunofluorescence Staining

The tissues, either paraffin-embedded or OCT-embedded, underwent sectioning with assistance from the Breast Center Pathology Core at Baylor College of Medicine. Paraffin-embedded slides were baked at 55°C for 2 hours, dewaxed, and rehydrated following standard protocols. Antigen retrieval was performed using Dako Retrieval Solution pH 9.0 (Agilent Technologies, S236784-2) in a pressure cooker at 115°C for 10 minutes. For frozen sections, slides were thawed at room temperature for 10 minutes and rinsed with PBS. All slides received treatment with 0.1M NH4Cl in PBS for 10 minutes to minimize autofluorescence and were subsequently blocked in 10% donkey serum in 0.4% Triton X-100 PBS for 1 hour at room temperature. In cases where mouse primary antibodies were used on mouse tissues, M.O.M. Blocking Reagent (Vector Laboratories, MKB-2213-1) was employed for additional blocking. The slides were then incubated with primary antibodies overnight at 4°C in 0.2% Triton X-100 PBS. Following three washes with PBS, slides were incubated with secondary antibodies for 1 hour at room temperature. Hoechst 33342(Thermo Fisher Scientific, 62249) was used for nuclear staining. After washing, slides were mounted with ProLongTM Gold antifade mountant(Invitrogen, P36934). Imaging was conducted using a LSM780 or LSM880(Zeiss) confocal microscope, Cytation 5 Epi-fluorescence microscope, and Zeiss Axioscan.Z1 scanner. Image analysis was performed using ZEN (Zeiss) or Fiji. Primary antibodies used in this study were as follows: anti-Biotin-APC(Miltenyi Biotec,130-113-288), anti-mRFP(Rockland,600-401-379), anti-GFP(Novus Biologicals,NB100-1614), anti-Granzyme B(Novus Biologicals,NB100-684), anti-F4/80(Cell Signaling technology,70076,for paraffin section), anti-F4/80(BIO-RAD,MCA497,for frozen section), anti-ERa(Thermo Fisher Scientific,MA1-80216), anti-CD8a(Thermo Fisher Scientific,14-0195-82), anti-ly6G(BioLegend,127602), anti-human CD68(Proteintech,25747-1-AP), anti-human Cytokeratin 8(DSHB,TROMA-I) and anti-human Cytokeratin 19(DSHB,TROMA-III).

#### *In vitro* macrophage differentiation and co-culture assay

Murine hindlimb bone marrow cells were isolated following the protocol outlined in a previous study^71^. Subsequently, the cells were cultured in DMEM medium for 5 hours to eliminate stromal cells. After this incubation period, unattached bone marrow cells were collected and reseeded into a new plate containing DMEM culture medium. These cells were then infected with the ERE-GFP-luciferase virus for 2 days. Following viral transduction, the bone marrow cells were differentiated into macrophages by the addition of 25 ng/ml recombinant murine M-CSF (Peprotech,315-02) for 3 days. Estrogen-stripped DMEM, consisting of phenol red-free DMEM (Thermo Fisher Scientific,21063029), 10% charcoal-stripped FBS (Sigma-Aldrich,F6765) and antibiotics, was used to culture for an additional 4 days, with the medium being changed every 2 days. Subsequently, 600,000 ERE-macrophages were cultured with or without 25,000 various types of cancer cells in a 12-well plate for 2-3 days. In the non-contact co--culture assay, we initially seeded 600,000 ERE-macrophages in the lower wells of a 12-well plate and incubated them for 30 minutes to promote cell attachment. Following this, 25,000 LLC1 cancer cells suspended in 200 μl of medium were added to the upper culture inserts (0.4-μm pore size, Greiner Bio-One,665640). The luminescence signal of the ERE-reporter was detected using an IVIS Lumina II (PerkinElmer) at the indicated time points, while the fluorescence signal of the ERE-reporter was acquired using an LSM880 confocal microscope.

#### *In vivo* reconstitution of ERE-macrophages

Mice were injected with LLC1 cells via intra-iliac artery (IIA) injection to induce bone metastatic tumors for a duration of 2 weeks. Following this, *in vivo* macrophage depletion was achieved by administering 200 μl clodronate-containing liposomes (Encapsula NanoSciences,CLD-8901) via retro-orbital injection for two consecutive days. Two days post-clodronate-liposome treatment, approximately 1,500,000 bone marrow-derived ERE-macrophages, as described above, were delivered into the mice via retro-orbital injection to reconstitute the macrophage-depleted mouse model. Four days thereafter, hindlimb bones were harvested for sectioning, and immunofluorescence staining was performed on the bone section slides. Imaging data were collected using an LSM880 confocal microscope.

#### mRNA Extraction and qRT-PCR

RNA extraction was performed using the Direct-zol RNA Miniprep Kit (Zymo Research, R2052) according to the manufacturer’s instructions. Subsequently, cDNA was synthesized from total RNA using the RevertAid First Strand cDNA Synthesis Kit (Thermo Fisher Scientific, K1622). For real-time PCR, PowerUp™ SYBR™ Green Master Mix (Thermo Fisher Scientific, A25741) was utilized on a CFX96 Real-Time PCR system (Biorad). The expression levels of GAPDH mRNA were employed as the internal control. All of the primer sequences provided in **key resource table**.

#### Protein Extraction and Western Blotting

For western blotting, bone marrow-derived macrophages were detached from the culture dish and lysed using RIPA buffer (10mM Tris-HCl, pH 8.0, 1mM EDTA, 0.5mM EGTA, 1% Triton X-100, 0.1% Sodium Deoxycholate, 0.1% SDS, 140mM NaCl). Protein concentration was determined by BCA assay (Pierce BCA Protein Assay Kit, Thermo Fisher Scientific, 23225). 25ng total protein were loaded onto Invitrogen NuPage® Novex® Gel (Thermo Fisher Scientific, NP0336BOX) for electrophoresis. Subsequently, proteins were transferred from the gel to the nitrocellulose membrane (Thermo Fisher Scientific, IB301002) using the iBlot™ Dry Blotting device (Invitrogen). Following a 1-hour blocking step with 5% milk, the membranes were incubated with anti-ERα (Thermo Fisher Scientific, MA1-80216), anti-ERβ (Santa Cruz Biotechnology, sc-390243), anti-PR (Santa Cruz Biotechnology, sc-7208) and anti-α Tubulin (Santa Cruz Biotechnology, sc-23948) antibodies overnight at 4°C with primary antibodies. The next day, membranes were probed with secondary HRP antibodies(Goat anti-Mouse IgG, Invitrogen, 31430)(Goat anti-Rabbit IgG, Invitrogen, 31460) and scanned using the Amersham Imager 600 system.

#### Data Preprocessing, Filtering, and Batch Correction

We utilized the cellranger pipeline for processing the multiplexed raw data. In each dataset and for every patient, we generated multiplexing and count assays based on CellRanger outputs, with all subsequent steps performed using Seurat packages (versions 4.0 and 5.0)^72^.

Specifically, sequencing files from TotalSeqC were demultiplexed through Seurat pipeline based on README table uploaded together with data.

Initially, within the Seurat framework, we conducted hashtag demultiplexing. To ensure data quality, we implemented a two-step doublet filtering process. Initially, we filtered cells based on demultiplexing results and hashtag assignments, retaining only the singlets (cells with a single barcode label). Subsequently, cells with fewer than 200 detected genes or displaying a deviation exceeding two-fold from the median gene count were excluded. Moreover, cells with more than 10% mitochondrial genes were systematically filtered out to minimize mitochondrial gene contamination.

We proceeded to normalize each individual dataset using the default parameters of the NormalizeData function. After normalization, the FindVariableFeatures function was employed with its default settings to identify variable genes within each normalized dataset. For data integration, we utilized the SelectIntegrationFeatures function, selecting the top 2,000 variable genes.

To identify anchors across all datasets, we compared different methods, including “rpca,” “cca,” and “harmony.” The “rpca” method was chosen due to its superior speed and efficiency^73^. These anchors were subsequently employed to integrate matrices using the IntegrateData function, specifying the parameter dims as 1:50.

#### Unsupervised Clustering and Cell Type Annotation

Within the integrated assay, we conducted scaling and PCA analysis. To visualize the data, we employed Uniform Manifold Approximation and Projection (UMAP) based on the top 50 principal components. To explore transcriptomic diversity, we utilized the FindClusters function with a range of resolutions from 0.2 to 5. We opted to label cell types using high-resolution clustering results to enhance annotation accuracy. We utilized machine learning support vector machines (SVM)^64–65^. We trained the SVM model using a well-annotated single-cell dataset integrated from various publicly available sources (GSE122464, GSE122465, GSE134098^54^, GSE128423^55^, GSE108885, GSE108891, GSE118436^56^, which collectively comprised over 500,000 cells from mice, covering more than 20 cell types and states. After predicting cell types in our dataset using the trained references, we conducted manual validation by assessing the expression of classical cell markers^74^.

#### Pseudo-bulk Transcriptomic Comparisons and Pathway Analysis

We employed pseudo-bulk analysis to determine differentially expressed genes (DEGs) with the goal of assessing cell type-specific expression disparities at the population level. The process encompassed several key steps: Firstly, we selected cells of interest corresponding to the desired cell type(s) as a prerequisite for the subsequent DE analysis. Subsequently, raw counts were obtained through quality control (QC) filtering applied to the designated cells earmarked for DE analysis. These counts, along with associated metadata, were integrated at the sample level. During the DE analysis, we used DESeq2 packages^75^ in R and the results were cross-validated through limma voom method. Following the identification of DEGs, we conducted Gene Set Enrichment Analysis (GSEA) using the ClusterProfiler R package^67^. The pathway dataset employed for this analysis was sourced from MSigDB (https://www.gsea-msigdb.org/gsea/msigdb/). To assess pathway activity across different cell types and biotin statuses, we utilized the AUCell R package^76^ to compute pathway activity scores. Visualization of these scores was achieved through violin plots using the AddModuleScore function. Significance statistics were calculated using the Wilcoxon rank-sum test.

#### Single-Cell Trajectory Analysis

We built a single-cell differentiation trajectory for macrophages across different biotin statuses using the scVelo tool^61^. The initial determination of macrophage subclusters was accomplished through an unsupervised method within Seurat. These subclusters exhibited distinct tissue contributions, suggesting potential variations in their differentiation trajectories. Notably, we already possessed prior knowledge indicating that macrophage cluster 5 (c5) represented osteoclasts, we are also aware that osteoclasts are differentiated from macrophages. Consequently, our trajectory inference primarily focused on these subclusters within the macrophage population.

#### Quantification and Statistical Analysis

The number of animals or independent replicates, along with the statistical tests used, is specified in the figure legends or figures. Preliminary experiments with small group sizes were conducted to determine appropriate group sizes for each *in vivo* experiment. Final results were consolidated from multiple batches of *in vivo* experiments, including preliminary ones. All biologically independent samples were included in statistical analyses. No animals reaching the experimental endpoint were excluded from the final analysis. Data quantification utilized GraphPad v10.1.2, with results presented as Mean ± SEM in bar and curve plots. Statistical methods are outlined in the figure legends, employing two-sided tests. Parametric statistics were applied to normally distributed data, while nonparametric statistics were utilized for comparing metastatic burden, such as bioluminescent intensities and derived data. A significance level of p < 0.05 was used to determine statistical significance. Further details are provided in the figure legend.

## REFERENCES

1. Ghajar, C.M., Peinado, H., Mori, H., Matei, I.R., Evason, K.J., Brazier, H., Almeida, D., Koller, A., Hajjar, K.A., Stainier, D.Y.R., et al. (2013). The perivascular niche regulates breast tumour dormancy. Nature Cell Biology 15, 807–817. 10.1038/ncb2767.

2. Price, T.T., Burness, M.L., Sivan, A., Warner, M.J., Cheng, R., Lee, C.H., Olivere, L., Comatas, K., Magnani, J., Kim Lyerly, H., et al. (2016). Dormant breast cancer micrometastases reside in specific bone marrow niches that regulate their transit to and from bone. Sci Transl Med 8, 340ra373. 10.1126/scitranslmed.aad4059.

3. Esposito, M., Mondal, N., Greco, T.M., Wei, Y., Spadazzi, C., Lin, S.C., Zheng, H., Cheung, C., Magnani, J.L., Lin, S.H., et al. (2019). Bone vascular niche E-selectin induces mesenchymal-epithelial transition and Wnt activation in cancer cells to promote bone metastasis. Nat Cell Biol 21, 627–639. 10.1038/s41556-019-0309-2.

4. Martin, J.D., Seano, G., and Jain, R.K. (2019). Normalizing Function of Tumor Vessels: Progress, Opportunities, and Challenges. Annu Rev Physiol 81, 505–534. 10.1146/annurev-physiol-020518-114700.

5. Wang, H., Tian, L., Goldstein, A., Liu, J., Lo, H.C., Sheng, K., Welte, T., Wong, S.T.C., Gugala, Z., Stossi, F., et al. (2017). Bone-in-culture array as a platform to model early-stage bone metastases and discover anti-metastasis therapies. Nat Commun 8, 15045. 10.1038/ncomms15045.

6. Wang, H., Tian, L., Liu, J., Goldstein, A., Bado, I., Zhang, W., Arenkiel, B.R., Li, Z., Yang, M., Du, S., et al. (2018). The Osteogenic Niche Is a Calcium Reservoir of Bone Micrometastases and Confers Unexpected Therapeutic Vulnerability. Cancer Cell 34, 823–839 e827. 10.1016/j.ccell.2018.10.002.

7. Wang, H., Yu, C., Gao, X., Welte, T., Muscarella, A.M., Tian, L., Zhao, H., Zhao, Z., Du, S., Tao, J., et al. (2015). The osteogenic niche promotes early-stage bone colonization of disseminated breast cancer cells. Cancer Cell 27, 193–210. 10.1016/j.ccell.2014.11.017.

8. Bado, I.L., Zhang, W., Hu, J., Xu, Z., Wang, H., Sarkar, P., Li, L., Wan, Y.W., Liu, J., Wu, W., et al. (2021). The bone microenvironment increases phenotypic plasticity of ER(+) breast cancer cells. Dev Cell 56, 1100–1117 e1109. 10.1016/j.devcel.2021.03.008.

9. Zhang, W., Bado, I.L., Hu, J., Wan, Y.W., Wu, L., Wang, H., Gao, Y., Jeong, H.H., Xu, Z., Hao, X., et al. (2021). The bone microenvironment invigorates metastatic seeds for further dissemination. Cell 184, 2471–2486.e2420. 10.1016/j.cell.2021.03.011.

10. Satcher, R.L., and Zhang, X.H. (2021). Evolving cancer-niche interactions and therapeutic targets during bone metastasis. Nat Rev Cancer. 10.1038/s41568-021-00406-5.

11. Mao, H., Hart, S.A., Schink, A., and Pollok, B.A. (2004). Sortase-mediated protein ligation: a new method for protein engineering. J Am Chem Soc 126, 2670–2671. 10.1021/ja039915e.

12. Ge, Y., Chen, L., Liu, S., Zhao, J., Zhang, H., and Chen, P.R. (2019). Enzyme-Mediated Intercellular Proximity Labeling for Detecting Cell-Cell Interactions. J Am Chem Soc 141, 1833–1837. 10.1021/jacs.8b10286.

13. Ombrato, L., Nolan, E., Kurelac, I., Mavousian, A., Bridgeman, V.L., Heinze, I., Chakravarty, P., Horswell, S., Gonzalez-Gualda, E., Matacchione, G., et al. (2019). Metastatic-niche labelling reveals parenchymal cells with stem features. Nature 572, 603–608. 10.1038/s41586-019-1487-6.

14. Lim, E., Metzger-Filho, O., and Winer, E.P. (2012). The natural history of hormone receptor-positive breast cancer. Oncology (Williston Park) 26, 688–694, 696.

15. Dees, E.C., and Carey, L.A. (2013). Improving endocrine therapy for breast cancer: it’s not that simple. J Clin Oncol 31, 171–173. 10.1200/JCO.2012.46.2655.

16. Nardone, A., De Angelis, C., Trivedi, M.V., Osborne, C.K., and Schiff, R. (2015). The changing role of ER in endocrine resistance. Breast 24 *Suppl 2*, S60–66. 10.1016/j.breast.2015.07.015.

17. Chakraborty, B., Byemerwa, J., Krebs, T., Lim, F., Chang, C.Y., and McDonnell, D.P. (2023). Estrogen Receptor Signaling in the Immune System. Endocr Rev 44, 117–141. 10.1210/endrev/bnac017.

18. Reed, S.A., Brzovic, D.A., Takasaki, S.S., Boyko, K.V., and Antos, J.M. (2020). Efficient Sortase-Mediated Ligation Using a Common C-Terminal Fusion Tag. Bioconjug Chem 31, 1463–1473. 10.1021/acs.bioconjchem.0c00156.

19. Fridy, P.C., Li, Y., Keegan, S., Thompson, M.K., Nudelman, I., Scheid, J.F., Oeffinger, M., Nussenzweig, M.C., Fenyo, D., Chait, B.T., and Rout, M.P. (2014). A robust pipeline for rapid production of versatile nanobody repertoires. Nat Methods 11, 1253–1260. 10.1038/nmeth.3170.

20. Rhee, J.M., Pirity, M.K., Lackan, C.S., Long, J.Z., Kondoh, G., Takeda, J., and Hadjantonakis, A.K. (2006). In vivo imaging and differential localization of lipid-modified GFP-variant fusions in embryonic stem cells and mice. Genesis 44, 202–218. 10.1002/dvg.20203.

21. Yu, C., Wang, H., Muscarella, A., Goldstein, A., Zeng, H.C., Bae, Y., Lee, B.H., and Zhang, X.H. (2016). Intra-iliac Artery Injection for Efficient and Selective Modeling of Microscopic Bone Metastasis. J Vis Exp. 10.3791/53982.

22. Zhang, W., Xu, Z., Hao, X., He, T., Li, J., Shen, Y., Liu, K., Gao, Y., Liu, J., Edwards, D.G., et al. (2023). Bone Metastasis Initiation Is Coupled with Bone Remodeling through Osteogenic Differentiation of NG2+ Cells. Cancer Discov 13, 474–495. 10.1158/2159-8290.CD-22-0220.

23. Sun, J., Hu, L., Bok, S., Yallowitz, A.R., Cung, M., McCormick, J., Zheng, L.J., Debnath, S., Niu, Y., Tan, A.Y., et al. (2023). A vertebral skeletal stem cell lineage driving metastasis. Nature 621, 602–609. 10.1038/s41586-023-06519-1.

24. Vanharanta, S., and Massague, J. (2013). Origins of metastatic traits. Cancer Cell 24, 410–421. 10.1016/j.ccr.2013.09.007.

25. Gao, Y., Bado, I., Wang, H., Zhang, W., Rosen, J.M., and Zhang, X.H. (2019). Metastasis Organotropism: Redefining the Congenial Soil. Dev Cell 49, 375–391. 10.1016/j.devcel.2019.04.012.

26. Li, X.F., Selli, C., Zhou, H.L., Cao, J., Wu, S., Ma, R.Y., Lu, Y., Zhang, C.B., Xun, B., Lam, A.D., et al. (2023). Macrophages promote anti-androgen resistance in prostate cancer bone disease. J Exp Med 220. 10.1084/jem.20221007.

27. Ma, R.Y., Zhang, H., Li, X.F., Zhang, C.B., Selli, C., Tagliavini, G., Lam, A.D., Prost, S., Sims, A.H., Hu, H.Y., et al. (2020). Monocyte-derived macrophages promote breast cancer bone metastasis outgrowth. J Exp Med 217. 10.1084/jem.20191820.

28. Hao, X., Shen, Y., Chen, N., Zhang, W., Valverde, E., Wu, L., Chan, H.L., Xu, Z., Yu, L., Gao, Y., et al. (2023). Osteoprogenitor-GMP crosstalk underpins solid tumor-induced systemic immunosuppression and persists after tumor removal. Cell Stem Cell 30, 648–664 e648. 10.1016/j.stem.2023.04.005.

29. Chakraborty, B., Byemerwa, J., Shepherd, J., Haines, C.N., Baldi, R., Gong, W., Liu, W., Mukherjee, D., Artham, S., Lim, F., et al. (2021). Inhibition of estrogen signaling in myeloid cells increases tumor immunity in melanoma. J Clin Invest 131. 10.1172/JCI151347.

30. Tanaka, Y., Nakayamada, S., and Okada, Y. (2005). Osteoblasts and osteoclasts in bone remodeling and inflammation. Current Drug Targets: Inflammation and Allergy. 10.2174/1568010054022015.

31. Xu, F., and Teitelbaum, S.L. (2013). Osteoclasts: New Insights. Bone Research. 10.4248/BR201301003.

32. Mass, E., Nimmerjahn, F., Kierdorf, K., and Schlitzer, A. (2023). Tissue-specific macrophages: how they develop and choreograph tissue biology. Nat Rev Immunol 23, 563–579. 10.1038/s41577-023-00848-y.

33. Coleman, R.E., Croucher, P.I., Padhani, A.R., Clezardin, P., Chow, E., Fallon, M., Guise, T., Colangeli, S., Capanna, R., and Costa, L. (2020). Bone metastases. Nat Rev Dis Primers 6, 83. 10.1038/s41572-020-00216-3.

34. Guise, T.A., Mohammad, K.S., Clines, G., Stebbins, E.G., Wong, D.H., Higgins, L.S., Vessella, R., Corey, E., Padalecki, S., Suva, L., and Chirgwin, J.M. (2006). Basic mechanisms responsible for osteolytic and osteoblastic bone metastases. Clin Cancer Res 12, 6213s–6216s. 10.1158/1078-0432.CCR-06-1007.

35. Kang, Y., Siegel, P.M., Shu, W., Drobnjak, M., Kakonen, S.M., Cordon-Cardo, C., Guise, T.A., and Massague, J. (2003). A multigenic program mediating breast cancer metastasis to bone. Cancer Cell 3, 537–549. 10.1016/s1535-6108(03)00132-6.

36. Weilbaecher, K.N., Guise, T.A., and McCauley, L.K. (2011). Cancer to bone: a fatal attraction. Nat Rev Cancer 11, 411–425. 10.1038/nrc3055.

37. Lee, J.W., Lee, I.H., Iimura, T., and Kong, S.W. (2021). Two macrophages, osteoclasts and microglia: from development to pleiotropy. Bone Res 9, 11. 10.1038/s41413-020-00134-w.

38. Clausen, B.E., Burkhardt, C., Reith, W., Renkawitz, R., and Forster, I. (1999). Conditional gene targeting in macrophages and granulocytes using LysMcre mice. Transgenic Res 8, 265–277. 10.1023/a:1008942828960.

39. Burgess, M., Wicks, K., Gardasevic, M., and Mace, K.A. (2019). Cx3CR1 Expression Identifies Distinct Macrophage Populations That Contribute Differentially to Inflammation and Repair. Immunohorizons 3, 262–273. 10.4049/immunohorizons.1900038.

40. Passegue, E., Wagner, E.F., and Weissman, I.L. (2004). JunB deficiency leads to a myeloproliferative disorder arising from hematopoietic stem cells. Cell 119, 431–443. 10.1016/j.cell.2004.10.010.

41. Qian, B.Z., and Pollard, J.W. (2010). Macrophage diversity enhances tumor progression and metastasis. Cell 141, 39–51. 10.1016/j.cell.2010.03.014.

42. Ruffell, B., and Coussens, L.M. (2015). Macrophages and therapeutic resistance in cancer. Cancer Cell 27, 462–472. 10.1016/j.ccell.2015.02.015.

43. de Visser, K.E., and Joyce, J.A. (2023). The evolving tumor microenvironment: From cancer initiation to metastatic outgrowth. Cancer Cell 41, 374–403. 10.1016/j.ccell.2023.02.016.

44. Sica, A., and Mantovani, A. (2012). Macrophage plasticity and polarization: in vivo veritas. J Clin Invest 122, 787–795. 10.1172/JCI59643.

45. Akkari, L., Bowman, R.L., Tessier, J., Klemm, F., Handgraaf, S.M., de Groot, M., Quail, D.F., Tillard, L., Gadiot, J., Huse, J.T., et al. (2020). Dynamic changes in glioma macrophage populations after radiotherapy reveal CSF-1R inhibition as a strategy to overcome resistance. Sci Transl Med 12. 10.1126/scitranslmed.aaw7843.

46. Mehta, A.K., Cheney, E.M., Hartl, C.A., Pantelidou, C., Oliwa, M., Castrillon, J.A., Lin, J.R., Hurst, K.E., de Oliveira Taveira, M., Johnson, N.T., et al. (2021). Targeting immunosuppressive macrophages overcomes PARP inhibitor resistance in BRCA1-associated triple-negative breast cancer. Nat Cancer 2, 66–82. 10.1038/s43018-020-00148-7.

47. Sun, L., Kees, T., Almeida, A.S., Liu, B., He, X.Y., Ng, D., Han, X., Spector, D.L., McNeish, I.A., Gimotty, P., et al. (2021). Activating a collaborative innate-adaptive immune response to control metastasis. Cancer Cell 39, 1361–1374 e1369. 10.1016/j.ccell.2021.08.005.

48. Cooke, P.S., Nanjappa, M.K., Ko, C., Prins, G.S., and Hess, R.A. (2017). Estrogens in Male Physiology. Physiol Rev 97, 995–1043. 10.1152/physrev.00018.2016.

49. Early Breast Cancer Trialists’ Collaborative, G. (2005). Effects of chemotherapy and hormonal therapy for early breast cancer on recurrence and 15-year survival: an overview of the randomised trials. Lancet 365, 1687–1717. 10.1016/S0140-6736(05)66544-0.

50. Jurga, A.M., Paleczna, M., and Kuter, K.Z. (2020). Overview of General and Discriminating Markers of Differential Microglia Phenotypes. Front Cell Neurosci 14, 198. 10.3389/fncel.2020.00198.

51. Bain, C.C., and MacDonald, A.S. (2022). The impact of the lung environment on macrophage development, activation and function: diversity in the face of adversity. Mucosal Immunol 15, 223–234. 10.1038/s41385-021-00480-w.

52. Wen, Y., Lambrecht, J., Ju, C., and Tacke, F. (2021). Hepatic macrophages in liver homeostasis and diseases-diversity, plasticity and therapeutic opportunities. Cell Mol Immunol 18, 45–56. 10.1038/s41423-020-00558-8.

53. Schulz, D., Severin, Y., Zanotelli, V.R.T., and Bodenmiller, B. (2019). In-Depth Characterization of Monocyte-Derived Macrophages using a Mass Cytometry-Based Phagocytosis Assay. Sci Rep 9, 1925. 10.1038/s41598-018-38127-9.

54. Baccin, C., Al-Sabah, J., Velten, L., Helbling, P.M., Grunschlager, F., Hernandez-Malmierca, P., Nombela-Arrieta, C., Steinmetz, L.M., Trumpp, A., and Haas, S. (2020). Combined single-cell and spatial transcriptomics reveal the molecular, cellular and spatial bone marrow niche organization. Nat Cell Biol 22, 38–48. 10.1038/s41556-019-0439-6.

55. Baryawno, N., Przybylski, D., Kowalczyk, M.S., Kfoury, Y., Severe, N., Gustafsson, K., Kokkaliaris, K.D., Mercier, F., Tabaka, M., Hofree, M., et al. (2019). A Cellular Taxonomy of the Bone Marrow Stroma in Homeostasis and Leukemia. Cell 177, 1915–1932 e1916. 10.1016/j.cell.2019.04.040.

56. Tikhonova, A.N., Dolgalev, I., Hu, H., Sivaraj, K.K., Hoxha, E., Cuesta-Dominguez, A., Pinho, S., Akhmetzyanova, I., Gao, J., Witkowski, M., et al. (2019). The bone marrow microenvironment at single-cell resolution. Nature 569, 222–228. 10.1038/s41586-019-1104-8.

57. Kim, I.S., Gao, Y., Welte, T., Wang, H., Liu, J., Janghorban, M., Sheng, K., Niu, Y., Goldstein, A., Zhao, N., et al. (2019). Immuno-subtyping of breast cancer reveals distinct myeloid cell profiles and immunotherapy resistance mechanisms. Nat Cell Biol 21, 1113–1126. 10.1038/s41556-019-0373-7.

58. Zheng, G.X., Terry, J.M., Belgrader, P., Ryvkin, P., Bent, Z.W., Wilson, R., Ziraldo, S.B., Wheeler, T.D., McDermott, G.P., Zhu, J., et al. (2017). Massively parallel digital transcriptional profiling of single cells. Nat Commun 8, 14049. 10.1038/ncomms14049.

59. Hao, Y., Stuart, T., Kowalski, M.H., Choudhary, S., Hoffman, P., Hartman, A., Srivastava, A., Molla, G., Madad, S., Fernandez-Granda, C., and Satija, R. (2024). Dictionary learning for integrative, multimodal and scalable single-cell analysis. Nat Biotechnol 42, 293–304. 10.1038/s41587-023-01767-y.

60. Wolf, F.A., Angerer, P., and Theis, F.J. (2018). SCANPY: large-scale single-cell gene expression data analysis. Genome Biol 19, 15. 10.1186/s13059-017-1382-0.

61. Bergen, V., Lange, M., Peidli, S., Wolf, F.A., and Theis, F.J. (2020). Generalizing RNA velocity to transient cell states through dynamical modeling. Nat Biotechnol 38, 1408–1414. 10.1038/s41587-020-0591-3.

62. La Manno, G., Soldatov, R., Zeisel, A., Braun, E., Hochgerner, H., Petukhov, V., Lidschreiber, K., Kastriti, M.E., Lonnerberg, P., Furlan, A., et al. (2018). RNA velocity of single cells. Nature 560, 494–498. 10.1038/s41586-018-0414-6.

63. Qiu, X., Zhang, Y., Martin-Rufino, J.D., Weng, C., Hosseinzadeh, S., Yang, D., Pogson, A.N., Hein, M.Y., Hoi Joseph Min, K., Wang, L., et al. (2022). Mapping transcriptomic vector fields of single cells. Cell 185, 690–711 e645. 10.1016/j.cell.2021.12.045.

64. Abdelaal, T., Michielsen, L., Cats, D., Hoogduin, D., Mei, H., Reinders, M.J.T., and Mahfouz, A. (2019). A comparison of automatic cell identification methods for single-cell RNA sequencing data. Genome Biol 20, 194. 10.1186/s13059-019-1795-z.

65. Alquicira-Hernandez, J., Sathe, A., Ji, H.P., Nguyen, Q., and Powell, J.E. (2019). scPred: accurate supervised method for cell-type classification from single-cell RNA-seq data. Genome Biol 20, 264. 10.1186/s13059-019-1862-5.

66. Piper, M., Mistry, M., Liu, J., Gammerdinger, W., and Khetani, R. (2022). hbctraining/scRNA-seq_online: scRNA-seq Lessons from HCBC (first release). In M. Piper, M. Mistry, J. Liu, W. Gammerdinger, and R. Khetani, eds. 1 ed.

67. Yu, G., Wang, L.G., Han, Y., and He, Q.Y. (2012). clusterProfiler: an R package for comparing biological themes among gene clusters. OMICS 16, 284–287. 10.1089/omi.2011.0118.

68. Hanzelmann, S., Castelo, R., and Guinney, J. (2013). GSVA: gene set variation analysis for microarray and RNA-seq data. BMC Bioinformatics 14, 7. 10.1186/1471-2105-14-7.

69. Hewitt, S.C., Kissling, G.E., Fieselman, K.E., Jayes, F.L., Gerrish, K.E., and Korach, K.S. (2010). Biological and biochemical consequences of global deletion of exon 3 from the ER alpha gene. FASEB J 24, 4660–4667. 10.1096/fj.10-163428.

70. Meerbrey, K.L., Hu, G., Kessler, J.D., Roarty, K., Li, M.Z., Fang, J.E., Herschkowitz, J.I., Burrows, A.E., Ciccia, A., Sun, T., et al. (2011). The pINDUCER lentiviral toolkit for inducible RNA interference in vitro and in vivo. Proc Natl Acad Sci U S A 108, 3665–3670. 10.1073/pnas.1019736108.

71. Nguyen, T., Du, J., and Li, Y.C. (2021). A protocol for macrophage depletion and reconstitution in a mouse model of sepsis. STAR Protoc 2, 101004. 10.1016/j.xpro.2021.101004.

72. Butler, A., Hoffman, P., Smibert, P., Papalexi, E., and Satija, R. (2018). Integrating single-cell transcriptomic data across different conditions, technologies, and species. Nat Biotechnol 36, 411–420. 10.1038/nbt.4096.

73. Luecken, M.D., Buttner, M., Chaichoompu, K., Danese, A., Interlandi, M., Mueller, M.F., Strobl, D.C., Zappia, L., Dugas, M., Colome-Tatche, M., and Theis, F.J. (2022). Benchmarking atlas-level data integration in single-cell genomics. Nat Methods 19, 41–50. 10.1038/s41592-021-01336-8.

74. Chen, J., Bardes, E.E., Aronow, B.J., and Jegga, A.G. (2009). ToppGene Suite for gene list enrichment analysis and candidate gene prioritization. Nucleic Acids Res 37, W305–311. 10.1093/nar/gkp427.

75. Love, M.I., Huber, W., and Anders, S. (2014). Moderated estimation of fold change and dispersion for RNA-seq data with DESeq2. Genome Biol 15, 550. 10.1186/s13059-014-0550-8.

76. Aibar, S., Gonzalez-Blas, C.B., Moerman, T., Huynh-Thu, V.A., Imrichova, H., Hulselmans, G., Rambow, F., Marine, J.C., Geurts, P., Aerts, J., et al. (2017). SCENIC: single-cell regulatory network inference and clustering. Nat Methods 14, 1083–1086. 10.1038/nmeth.4463.

